# Impact of adolescent high-fat diet and psychosocial stress on neuroendocrine stress responses and binge eating behavior in adult male Lewis rats

**DOI:** 10.1101/2024.11.12.623254

**Authors:** Julio Sierra, Timothy B. Simon, Darine Abu Hilal, Yaria Arroyo Torres, José M. Santiago Santana, Johnny D. Figueroa

## Abstract

Childhood obesity is a multifactorial disease affecting more than 160 million adolescents worldwide. Adolescent exposure to obesogenic environments, characterized by access to high-fat diets and stress, precipitates maladaptive eating habits in adulthood such as binge eating. Evidence suggests a strong association between Western-like high-saturated-fat (WD) food consumption and dysregulated hormone fluctuations. However, few studies have explored the long-term impact of adolescent WD and psychosocial stress on brain and behavior. This longitudinal study aimed to investigate the impact of adolescent exposure to an obesogenic diet on stress resiliency and increased susceptibility for binge-like eating behaviors. Adolescent male Lewis rats were given WD (41% fat; n=40) or control diet (CD, 16% fat; n=38) for 4 weeks before undergoing a stress paradigm of predator exposure and social instability (CDE, WDE, CDU, WDU; n=16/group). Subjects were provided intermittent WD access (24 h/week) to evaluate binge eating–like behavior in adulthood. Fecal corticosterone and testosterone were measured at four timepoints throughout adolescence and adulthood. WD rats exhibited increased body weight (p = 0.0217) and elevated testosterone in mid-adolescence (p=0.0312) and blunted stress-induced corticosterone response in mid-late adolescence (CDE:WDE, p=0.028). Adolescent hormone levels were negatively correlated with bingeing and explained the variability between adult rats expressing hyperphagic and hypophagic behaviors. These results demonstrate that exposure to WD in adolescence disrupts hormone fluctuations and stress responsivity, with effects persisting into adulthood. This underscores the importance of addressing obesogenic environments early to mitigate their lasting impact on hormone regulation and stress responsiveness.

## INTRODUCTION

Obesity constitutes a serious health issue affecting more than 160 million adolescents worldwide, with rates quadrupling in the last three decades (Phelps et al., 2024). While obesity is a multifactorial disease, increased access to Western-like diets enriched in saturated fats is a main contributor to the global obesity epidemic (Leigh et al., 2018). Acute and chronic (i.e., continuous, intermittent) consumption of Western high-saturated-fat diets (WD) during adolescence have been shown to impair metabolic and cognitive function (Boitard et al., 2014; Kaakoush et al., 2017; Kendig et al., 2021). However, additional research is needed to clarify the biological pathways during adolescence that influence Western diet intake and its long-term adverse effects on brain function and behavior.

Psychological stress is a powerful contributor to eating alterations and excessive weight gain, and prolonged or repeated exposure to stressors results in behavioral, biochemical, and physiological changes implicated in obesity (Tomiyama, 2019). Several clinical studies have reported a positive association between perceived stress and changes in dietary patterns (i.e., higher intake of saturated fat) (Laugero et al., 2011; Michels et al., 2013). Emotional eating reduces feelings of stress, but habitual consumption of “comfort food” results in abdominal obesity (Dallman, 2010). Evidence supports that elevated glucocorticoid levels drive palatable food intake to reduce central stress response activity (Dallman et al., 2005; Foster et al., 2009). Despite this, findings are sparse regarding the long-term effects of diet composition on stress reactivity (Jakulj et al., 2007; Shively et al., 2023).

Adolescence is a crucial period for brain development, particularly for the neural systems that regulate stress responsivity, making the brain highly susceptible to external influences such as stress. The hypothalamic–pituitary–adrenal (HPA) and hypothalamic–pituitary–gonadal (HPG) axes are vital neuroendocrine systems that mature during adolescence and play essential roles in modulating stress responses, maintaining homeostasis, and regulating puberty through the release of steroid hormones (Marceau et al., 2014). However, the maturation of these systems can be disrupted by environmental factors such as diet and stress, potentially leading to neuroinflammation and hormone signaling dysregulation. For example, studies in rodents demonstrate that high-saturated-fat diets induce inflammation in key brain regions associated with stress regulation, such as the hypothalamus, hippocampus, and amygdala (Boukouvalas et al., 2010; Buwalda et al., 2001; Santana et al., 2021; Tamashiro et al., 2006; Wang et al., 2022). This vulnerability to external influences is further compounded by the emergence of sex-dependent differences in hormonal responses during the peri-pubertal phase, shaping behavioral outcomes and stress resilience (Pascoe et al., 1991; Tannenbaum et al., 1997; Toniazzo et al., 2018). For instance, males exhibit higher cortisol reactivity than females in response to psychosocial stress (Stephens et al., 2016). Higher levels of testosterone are associated with decreased risk for disordered eating in males (Culbert et al., 2014). Thus, understanding how these systems interact with environmental stressors is critical for elucidating the neurobiological mechanisms underlying behavioral maturation and vulnerability to stress- related disorders.

The current longitudinal study investigated the impact of adolescent access to an obesogenic diet on compulsive eating behavior during adulthood. We hypothesized that early consumption of a Western-like, highly palatable, high-saturated-fat diet followed by psychosocial stress would result in maladaptive, compulsive eating behaviors in adulthood. We found that continuous high-fat diet consumption in adolescence induces an abnormal neuroendocrine stress response that persists in adulthood. This study enhances our knowledge of the neuroendocrine changes that may underpin dysregulated stress responses that promote maladaptive eating behaviors.

## MATERIALS AND METHODS

### Animals

Lewis rats are highly susceptible to environmental stressors and inflammatory challenges due to attenuated HPA axis activity (Stöhr et al., 2000). Thus, Lewis rats provide a suitable model for investigating the effects of obesogenic environments on a vulnerable population. Focusing exclusively on male rats can help isolate neuroendocrine mechanisms involved in stress-induced behavioral changes, enhancing our insights into the role of male-specific hormonal responses in the context of diet and stress. The decision to use only males eliminates potential variability due to sex differences in hormone concentrations during pubertal development. This simplifies data analysis and interpretation, while maintaining consistency with previous research conducted by our team.

Experimental procedures were conducted under the approval of the Institutional Animal Care and Use Committee (IACUC) at Loma Linda University. This study follows the ARRIVE 2.0 guidelines for reporting animal research (Sert et al., 2020). Female Lewis rat dams with male pups (postnatal day 15, PND15) were obtained from Charles River Laboratories (Portage, MI, USA). Upon arrival, female dams were housed with their pups and given *ad libitum* access to food and water. All rats were assessed to ensure they were healthy and that no adverse conditions were present. Animals were kept in standard housing conditions (12-hr light/dark cycle with lights on at 7:00 AM, 21 ± 2°C, and relative humidity of 30%) and allowed to acclimatize to the facility for one week before the start of the experiment. Adolescent male pups (PND21) were weaned, matched across diet groups by body weight, and pair-housed with *ad libitum* access to assigned diets and water for the duration of the study. During the weaning procedure, one pup was found deceased, so the matched cage partner was removed from the study.

### Study Design

Adolescent Lewis rats (PND21) were weight-matched and randomized to receive either a Western-like high-saturated-fat diet (WD, n=40) or an ingredient-matched purified control diet (CD, n=38). Animals on the WD were given 60 grams of low-fat diet to provide the option to consume either diet. The WD (*Product No. F7462*; 41.4% kcal from fat) and CD (Product No. F7463; 3.8 kcal/gram, 16.5% kcal from fat) were obtained from Bio-Serv (Frenchtown, NJ, USA). The macronutrient composition and fatty acid profiles are detailed in previous studies and summarized in **Supplemental Table 1** (Vega-Torres et al., 2022, 2020). Rats were allowed to consume their respective diets for 4 weeks. A subset of rats (PND51) from each group were euthanized and brain tissue was collected for further analysis (n=6-8/group). The remaining subjects were further subdivided into one of four groups: **(1)** control diet, unexposed (CDU); **(2)** control diet, exposed (CDE); **(3)** Western diet, unexposed (WDU); and **(4)** Western diet, exposed (WDE; n=16/group). Subjects in the exposed subgroups underwent a protocol of psychosocial stress (PSS), consisting of one exposure to a live predator followed by 10 days of social instability (PND54–64). Following the end of PSS, all groups were introduced to a binge eating paradigm, consisting of an initial 48-hour exposure to the WD, followed by weekly 24-hour re-exposure to the WD for three weeks. One day before the end of the experiment, all animals were given WD for 24 hours. Subjects were euthanized, and plasma and brain tissue were collected. The experimental timeline is illustrated in **Figure 1**.

**Figure 1.**
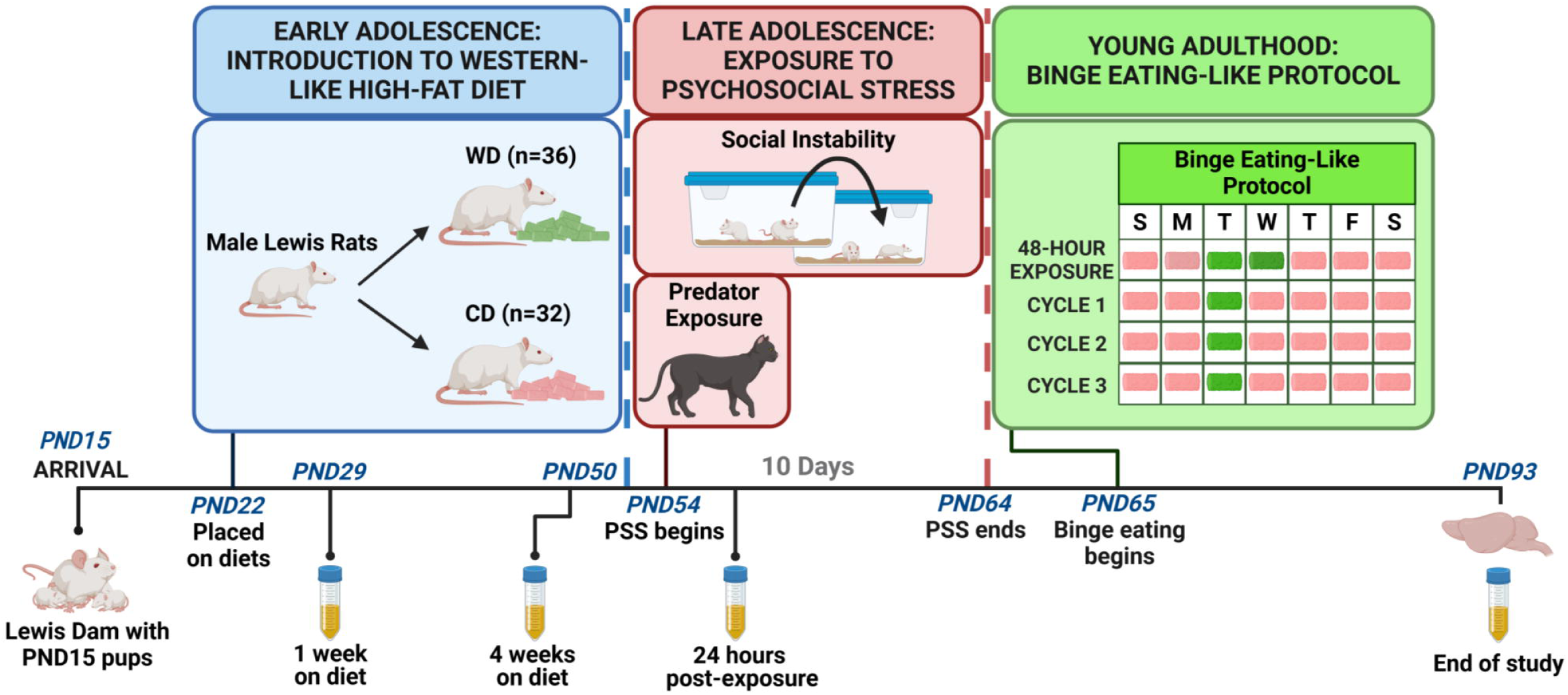
Experimental timeline. Rat pups (PND15) arrived and were allowed to acclimate for one week. Adolescent rats were weight-matched and separated into one of two groups: control diet (CD, n=38) or Western-like diet (WD, n=44). After four weeks on their respective diets, animals were further subdivided into one of four groups: control diet unexposed (CDU), control diet exposed (CDE), Western diet unexposed (WDU), and Western diet exposed (WDE) (n=16-20/group). Late adolescent rats (PND50) in the exposed subgroups underwent psychosocial stress exposure consisting of a one-hour exposure to a cat followed by 10 days of social instability. Following stress exposure, all animals (PND60) were introduced to a four-week binge-eating protocol (BED). During the first week, rats were provided with WD for 48 hours. After 48 hours, WD was replaced with CD for the remainder of the week. This was repeated for three more cycles, but WD access was limited to 24 hours. All animals were euthanized on PND92.

Weekly bodyweight and food consumption measurements were assessed manually (0800h-1000h). Food intake was measured in grams by calculating the difference between the total chow provided and the remaining chow at the next feeding. Food intake measurements were divided in half to account for pair-house conditions. Energy intake (kcal) was derived from food intake values using macronutrient information (CD, 3.8 kcal/gram; WD, 4.6 kcal/g).

### Psychosocial Stress Model

To evaluate whether access to an adolescent high-saturated-fat diet impacts susceptibility to perceived stressors during young adulthood, we adapted an established rat model of traumatic stress, consisting of trauma-inducing exposure to a natural predator and social instability (Ontiveros-Ángel et al., 2024; Sharafeddin et al., 2024; Zoladz et al., 2012).

#### Predator exposure

The exposure was performed for one hour during the light cycle (8:00–9:00am). We employed the use of a domesticated mature female cat, including the collection of soiled cat litter, sifted for stool. Rats were moved to the testing area one hour before the start of the experiment and immobilized using DecapiCones (Cat. No. NC9679094, Braintree Scientific; MA, USA). The restrained animals were placed in a perforated wedge-shaped plexiglass pie cage designed for aerosol delivery (Cat. No. RPC-1 AERO, Braintree Scientific; MA, USA; Diameter 41 cm × Height 6.75 cm). The plexiglass cage was placed inside a larger metal enclosure (91.4 cm × 58.4 cm × 63.5 cm, Amazon Basics, Amazon, USA) and connected to a nebulizer to deliver aerosolized cat litter odor to the animals. Then, the cat was brought into the testing room and placed in the metal enclosure. To minimize variability, personnel were not allowed in the room during the exposure. Animals were continuously observed by video call for the duration of the experiment. Subjects were returned to their cages and monitored for 30 minutes immediately after exposure, with an additional check-in the following day.

#### Social instability

Subjects were exposed to unstable housing conditions for 10 days starting the day after predator exposure. Cage partners were swapped daily at a random time each day within their respective subgroups to prevent cross-diet exposure. Subjects were not housed with the same cage partner on successive days. All animals were returned to their original partner on the final day of the social instability protocol. Unexposed rats were undisturbed and remained housed with the same cage mate for the duration of the study.

### Binge Eating–Like Paradigm

All animals, irrespective of group, underwent a four–week experimental paradigm to induce bingeing behaviors (Czyzyk et al., 2010). This model can induce binging behavior within one week and does not require food restriction or stress exposure. During the initial week, animals were provided with the high-saturated-fat WD and the low-fat CD for 48 hours. After 48 hours, the WD was removed and replaced with the CD for the remaining five days. The following week, animals received WD and CD for 24 hours. Food consumption measurements were taken at 2.5- and 24-hours. After 24 hours, the WD was removed and replaced with the CD for the rest of the week (cycle 1). This was repeated for two additional weeks (i.e. cycles 2/3). The WD was provided on the same day of the week during the light cycle each week.

### Tissue Collection

Subjects were deeply anesthetized with 3-5% isoflurane before undergoing transcardial perfusion with 0.01 M phosphate-buffered saline (PBS), prepared beforehand and pre-chilled at 4°C. Subjects were euthanized and brain tissue was collected. All tissue was immediately submerged in RNAlater™ Stabilization Solution (Cat. No. AM7021; Thermofisher, Waltham, MA, USA). Samples were stored at -80°C until further processing.

### Western Immunoblotting

Unilateral (left hemisphere) micropunches were obtained from the hippocampus. The tissue was transferred to a 1.5 mL micro tube containing 700 µl of cold extraction buffer (Cell Lytic MT Lysis Extraction Buffer, cat. no. C3228, Sigma-Aldrich, St. Louis, MO, USA), 1% phosphatase inhibitor cocktail 3 (cat. no. P0044, Sigma-Aldrich, St. Louis, MO, USA), and Sigma FAST Protease Inhibitor Cocktail Tablet, EDTA free (cat. no. S8830, Sigma-Aldrich, St. Louis, MO, USA). The samples were treated according to the Lysis Buffer manufacturer’s suggested directions. The samples were centrifuged (20,000 rpm) for 10 minutes at 4°C and the supernatant was collected and stored at -80°C until needed for further processing. Protein quantification was performed using the Bio-Rad protein assay according to the manufacturer’s instructions (Bio-Rad Laboratories, Hercules, CA, USA). The proteins were then separated on a 10% polyacrylamide-SDS gel (90 µg of protein/lane) and wet transferred to a nitrocellulose membrane for 1h at 4°C. The membrane was blocked with Odyssey Blocking Buffer (cat. no. 927-40000, LI-COR Biosciences, Lincoln, NE, USA) for 1 hour at room temperature. Immunodetection was done by adding the primary antibody, rabbit IL-6R alpha polyclonal antibody (1:500; cat. no. 23457-1-AP, Proteintech, Rosemont, IL, USA), in blocking solution and incubating overnight at 4°C. Anti-mouse ß-actin (1:5000; cat. no. A5441, Sigma-Aldrich, St. Louis, MO, USA) was used as the loading control. For secondary antibodies, we used goat anti-rabbit (1:25000; cat. no. 926-32211, LI-COR Biosciences, Lincoln, NE, USA) and goat anti-mouse (1:25000; cat. no. 926-68070, LI-COR Biosciences, Lincoln, NE. USA) for 1h at room temperature. Infrared signals from membranes were detected using the LICOR Odyssey, model CLx Scanner (LI-COR Biosciences, Lincoln, NE, USA). Immunoblot densitometry analyses were quantified with Image Studio 5.2 Software (LI-COR Biosciences, Lincoln, NE, USA).

### Fecal Sample Collection and Metabolite Extraction

In order to determine corticosterone and testosterone concentrations, fecal metabolite extraction was performed as previously described (Kalyan-Masih et al., 2016; Vega-Torres et al., 2019, 2018). Fecal samples were collected at the following timepoints: **1)** one week after introduction to diets, **2)** four weeks after introduction to diets, **3)** 24 hours post-predator exposure, and **4)** at the end of the study, corresponding to early adolescence, mid-late adolescence, late adolescence, and young adulthood, respectively (Spear, 2000). Cages were changed 24 h before each collection. Fecal boli were collected and stored at -80°C until further processing. Samples were defrosted and air dried at room temperature for 30 minutes before weighing. Samples (1 ± 0.05 g) were manually pulverized, suspended in 5 mL of 70% ethanol, and placed on a rotator overnight (14-18 hours). The following day, samples were centrifuged (1363 x g, RCF) for 15 minutes at 4°C and the supernatant was transferred to clean tubes. Samples were centrifuged once more and transferred to clean tubes to ensure complete removal of solid fecal content.

### Fecal metabolite analysis

Steroid metabolite levels were evaluated by colorimetric competitive enzyme immunoassay kits. Fecal corticosterone and fecal testosterone concentrations were measured using Enzo Corticosterone ELISA Kit (sensitivity: 27.0 pg/mL; range: 32–20,000 pg/mL) (cat. no. ADI-900-097, Enzo Life Sciences, Farmingdale, NY, USA), and Arbor Assays DetectX Testosterone ELISA Kit (sensitivity: 9.92 pg/mL; range: 40.96–10,000 pg/mL) (cat. no. K032- H1W, Arbor Assays, Ann Arbor, MI, USA) respectively, according to the manufacturers’ instructions. Extracted fecal samples were diluted in kit assay buffer (1:20 dilution). Plate absorbance was read at 405 nm for corticosterone and 450 nm for testosterone, with a 570 nm correction for both, using the SpectraMax i3X detection platform (Molecular Devices, Sunnyvale, CA). Steroid concentration was back calculated by interpolation using a 4 Parameter Logistic Curve (4PLC) fit. Interpolated values were corrected by the dilution factor (20x). Inter- and intra-assay coefficient of variation (%CVs) were <15%.

### Statistical Analysis

All data were analyzed using GraphPad Prism 10 (GraphPad Software, La Jolla, CA, USA). Two-way ANOVA was used to examine the effect of diet, stress, and interaction between factors on outcome measures. Post hoc analyses were conducted using Tukey’s, Dunn’s (following Kruskal-Wallis test), Dunnett’s (following Welch’s ANOVA), or Sidak’s (following repeated measures two-way ANOVA) tests. Adjusted p-values were used in the case of multiple comparisons. We report eta-squared (η^2^) values as the measure of effect size, if applicable. Pearson correlation and principal component analysis (PCA) was used to evaluate the relationship between molecular targets and behavioral outcomes. Normality and equality of variance were assessed using the Shapiro-Wilk and Brown-Forsythe tests, respectively. The ROUT method was used to investigate outliers. Differences were considered significant for p < 0.05. All data are shown as the mean ± SEM.

## RESULTS

### Western Diet Exposure During Adolescence Increases Body Weight and Caloric Intake in Male Rats

To investigate the longitudinal effects of adolescent high-fat diet consumption on stress reactivity and eating behavior, we recorded weekly body weight **(Figure 2A)** and weight-corrected food consumption **(S Figure 1)**. Although diets were ingredient-matched, the proportion of macronutrients and caloric content varies between diets **(S Table 1)**. Therefore, we calculated energy intake (kcal/g) to provide additional insight into changes in eating behavior **(Figure 2B)**. WD animals displayed greater caloric intake (p < 0.0001) compared to the CD group in the first week (p < 0.0001). Individuals consuming WD exhibited increased body weight compared to controls after three weeks on the diet (p = 0.0217). Repeated measures two-way ANOVA revealed significant diet [F_(1, 80)_ = 5.886, p = 0.0175] and time [F_(1.365, 109.2)_ = 18165, p < 0.0001] main effects and diet x time interaction [F_(4, 320)_ = 19.58, p < 0.0001] effect on body weight. Statistics for weekly body weight, food consumption, and caloric intake are shown in **Table S2-4**, respectively.

**Figure 2.**
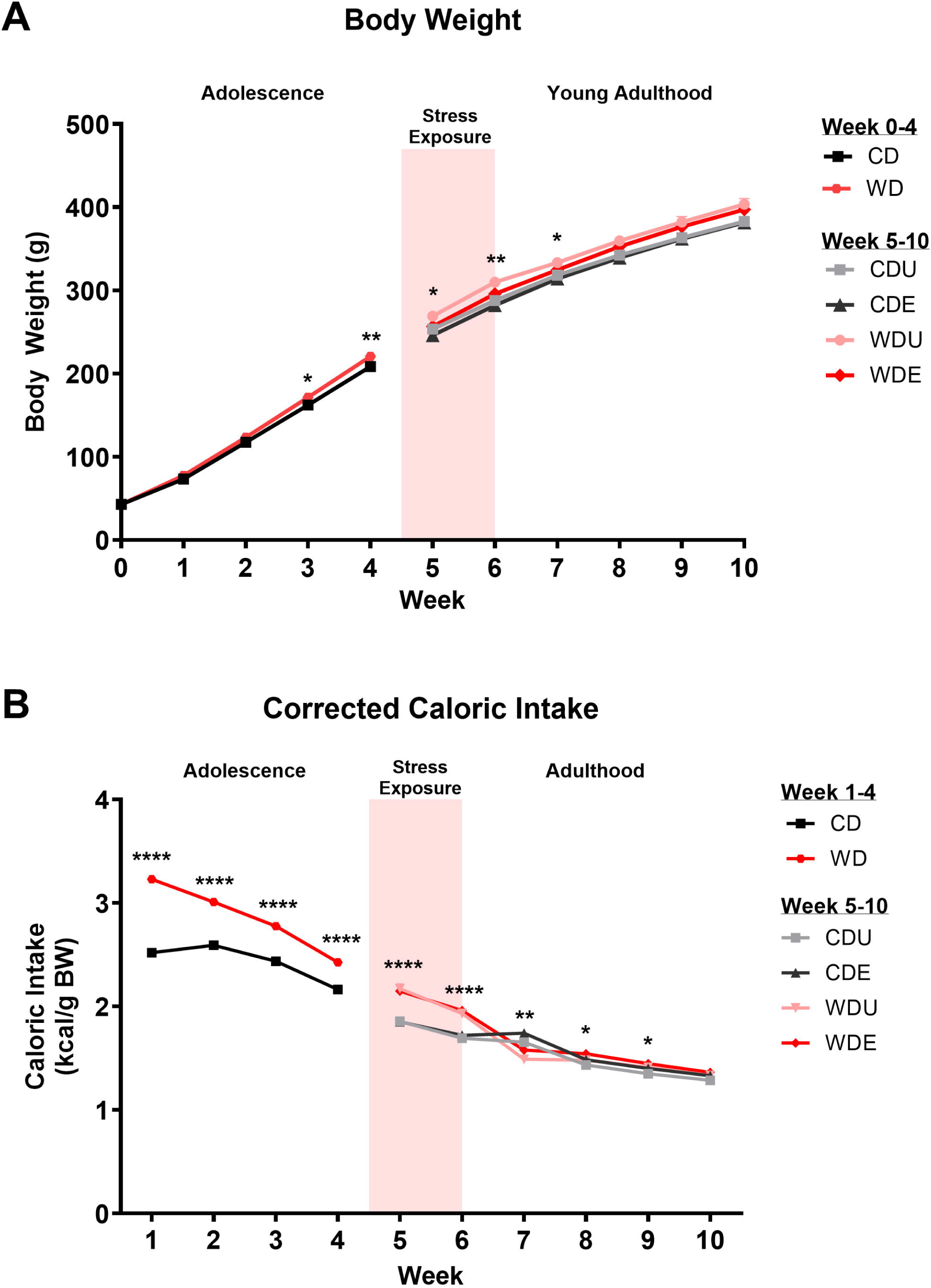
Western diet during adolescence increases body weight and caloric intake. Weekly average body weight and weight-corrected caloric intake for control diet (CD) and Western diet (WD) groups. Groups were subdivided into PSS-exposed (E) and unexposed (U) subgroups after four weeks. **A)** WD consumption led to an increase in body weight after three weeks on the diet compared to controls (p = 0.0120), which were maintained until the seventh week. **B)** WD animals displayed increased caloric intake until the final week of the study. Sample size n=32/group. * *p*<0.05, ** *p*<0.01, *** *p*<0.001

### Psychosocial Stress Exposure During Adolescence Enhances Body Weight Gain in Male Rats Consuming an Obesogenic Western Diet

Previous work from our lab has shown that consuming an obesogenic diet during adolescence alters neural and behavioral markers associated with emotional regulation in adults, with chronic psychosocial stress exacerbating these impacts (Kalyan-Masih et al., 2016; Ontiveros-Ángel et al., 2024; Sharafeddin et al., 2024; Vega-Torres et al., 2018). Building upon previous investigations, we sought to determine the impact of adolescent WD consumption on susceptibility to acute and chronic stress. To address this, we modified the psychosocial stress protocol to specifically target the transition period from adolescence to adulthood (Spear, 2000). Psychosocial stressors were introduced during late adolescence (PND54) with a single predator exposure (severe acute stress) followed by ten days of social instability (chronic stress).

There was a significant effect of stress [F_(1,58)_ = 7.747, p = 0.0072] on body weight change after predator stress. However, post-hoc analyses did not reveal any group differences **(Figure 3A)**. WDE rats displayed reduced food intake following acute stress compared to the WDU group (p = 0.0030) with a significant main effect of stress [F_(1,60)_ = 13.36, p = 0.0005] **(S Figure 2)**. A similar decrease was not seen in the CDE group compared to the CDU group. No differences were observed in weight-adjusted food consumption after predator exposure. Relative change in consumption from before psychosocial stress exposure was influenced by stress [F_(1,60)_ = 9.291, p = 0.0034] and diet [F_(1,60)_ = 10.05, p = 0.0024] **(Figure 3B)**. CDE animals showed decreased change in food intake compared to WDU animals (p = 0.0003) after severe acute stress.

**Figure 3.**
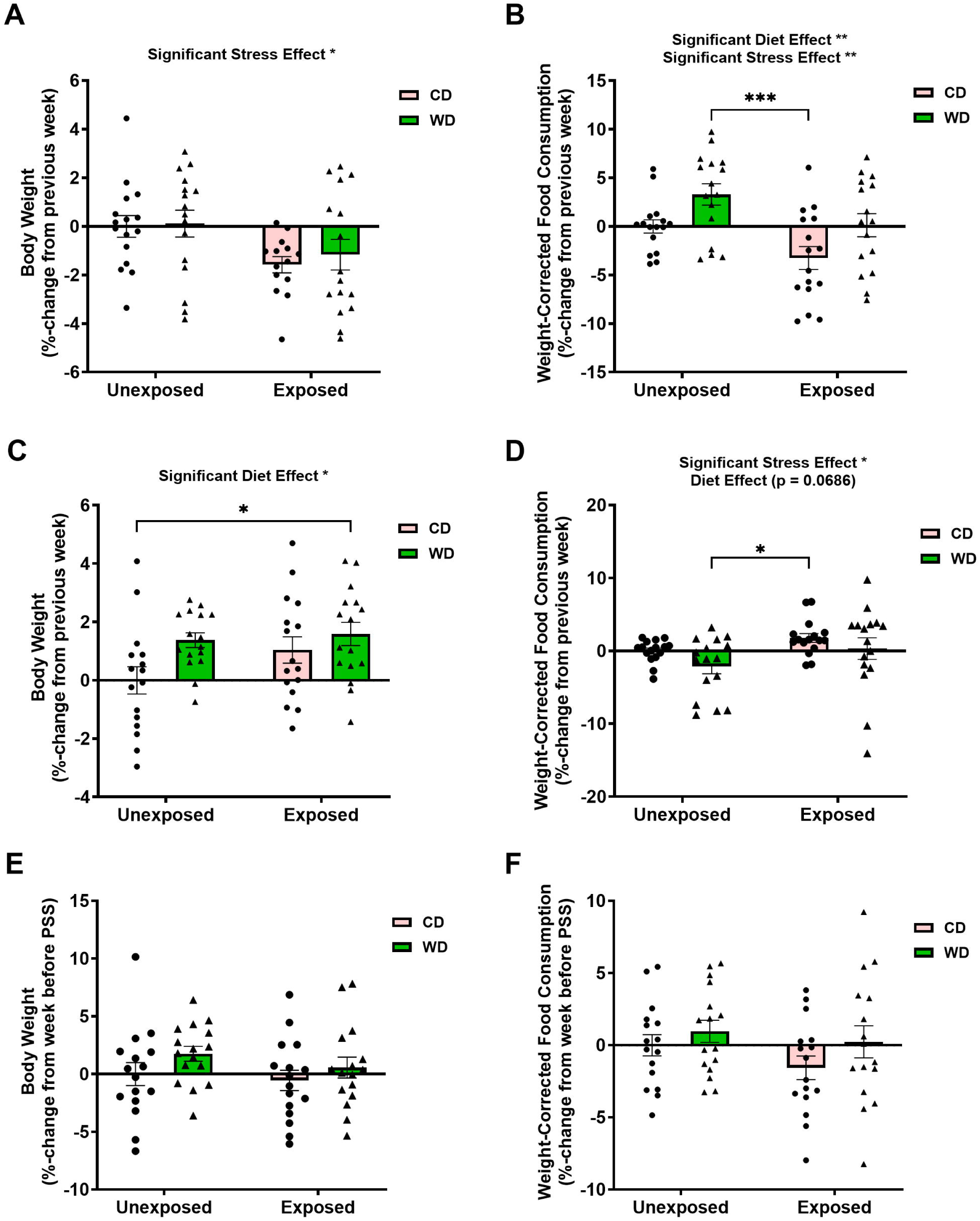
Psychosocial stress exposure enhances weight gain in male rats consuming an obesogenic Western-like diet during adolescence. **A)** Acute exposure to a live predator revealed a significant main effect of stress on body weight [F_(1,58)_ = 7.747, p = 0.0072]. Predator stress resulted in a trending decrease in body weight gain (CDU:CDE, p = 0.0723) **B)** There was a significant effect of stress [F_(1,60)_ = 9.291, p = 0.0034] and diet [F_(1,60)_ = 10.05, p = 0.0024] on decreasing and increasing food intake, respectively. **C)** WDE rats undergoing social instability exhibited higher weight gain than controls (WDE:CDU, p = 0.0273). **D)** Social instability stress [F_(1,60)_ = 4.798, p = 0.0324] impacted food intake in CD compared to unexposed WD animals (CDE:WDU, p = 0.0289). **E,F)** There were no differences in body weight or food intake due to predator and social instability stressors from before PSS introduction. n=16/group. * *p*<0.05, ** *p*<0.01, *** *p*<0.001

Following social instability, WDE rats exhibited greater body weight gain compared to controls (p < 0.0273), with a significant diet effect [F_(1,60)_ = 5.725, p = 0.0199] **(Figure 3C).** There was a diet effect [F_(1,60)_ = 17.54, p < 0.0001] on weight-adjusted food intake **(S Figure 3)**. CD animals experiencing social instability displayed a relative increase in consumption compared to unexposed WD animals (p = 0.0289), with a significant influence of stress [F_(1,60)_ = 4.798, p = 0.0324] and trend-like significance of diet [F_(1,60)_ = 3.438, p = 0.0686] **(Figure 3D)**. No differences were observed in body weight or food consumption change between groups across the entire PSS protocol from the week before PSS **(Figure 2E-F)**. There was a significant main effect of diet [F_(1,60)_ = 11.12, p = 0.0015] on total food consumed **(S Figure 4)**.

### Intermittent Access to the Western Diet Leads to Bingeing Behaviors in Rats Exposed to Psychosocial Stress During Adolescence

We adapted a protocol of cyclic (re-)exposure to a palatable high-fat diet that avoids the use of food restriction to examine bingeing behaviors (Czyzyk et al., 2010; Sharafeddin et al., 2024). One criterion for binge eating is the consumption of an abnormally large quantity of food in a short period of time (American Psychiatric Association, 2022). To determine whether diet and/or stress exposure induced a bingeing phenotype in our rats, we measured food consumption at 2.5 and 24 hours after re-introduction to the WD each week **(Figure 4)**. There were no differences in 2.5-hour or 24-hour corrected food consumption during the first or third binge-eating cycles. During the second BE cycle, stress exposure [F_(1,58)_ = 4.809, p = 0.0323] led to a decrease in 2.5-hour consumption. Further analysis revealed a trending difference between WD groups (p = 0.0661) **(Figure 4B)**. Interestingly, the WDE group showed increased food consumption compared to controls during the first (CDU:WDE, p = 0.0061) and third (CDU:WDE, p = 0.0157) BE cycles in the 24 hours following re-introduction to the control diet **(S Figure 5)**. WDE animals demonstrated an increased weekly food intake during the first (CDU:WDE, p =0.0021) and second (CDU:WDE, p = 0.0114), but not the third BE cycle **(S Figure 6)**.

**Figure 4.**
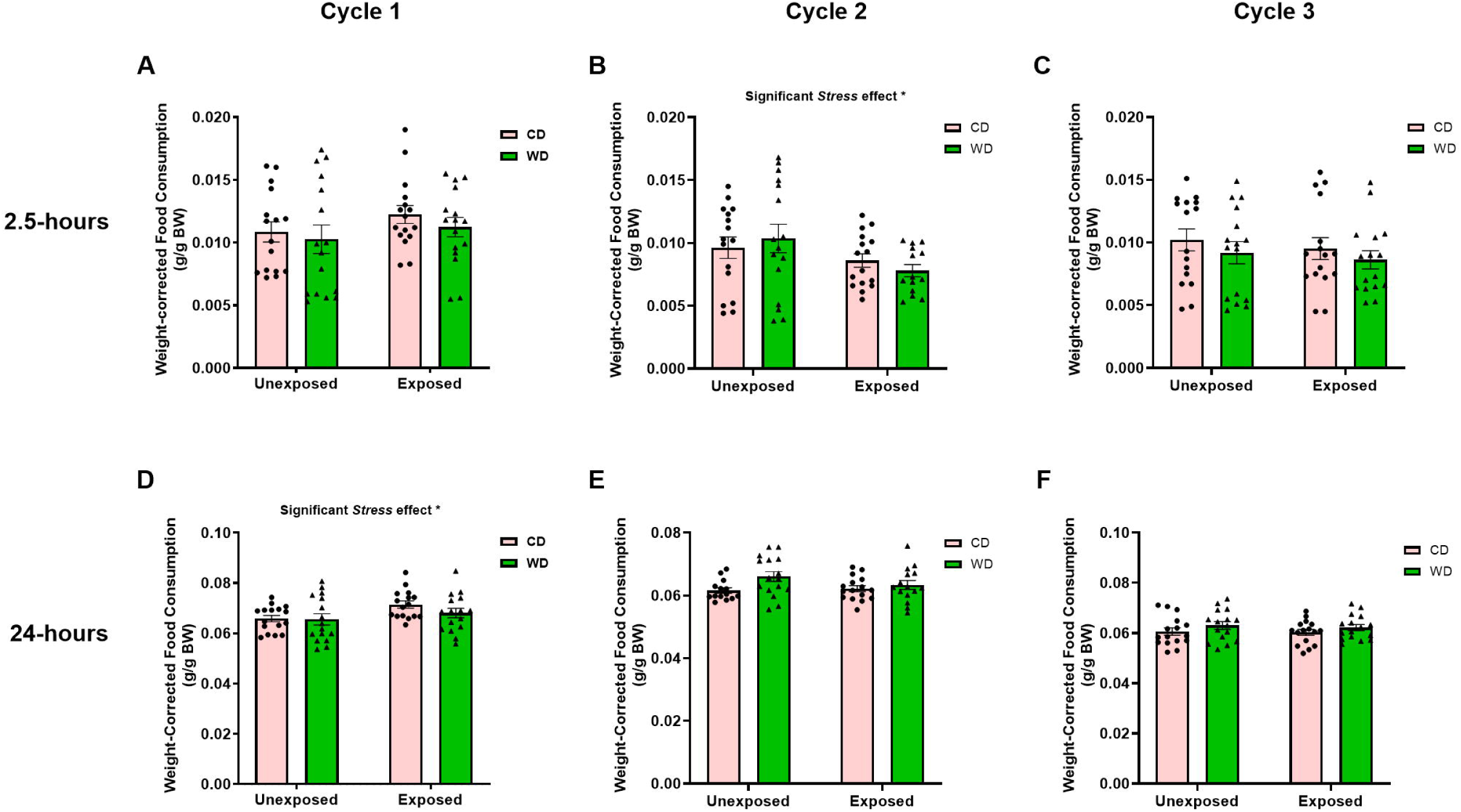
Intermittent WD access leads to binging behaviors in rats exposed to adolescent psychosocial stress. Food consumption was measured at 2.5 and 24 hours after weekly re-introduction to the WD (cycle 1-3). **A,C)** Analysis revealed no differences at 2.5-hour consumption during the first or third binge eating cycles. **B)** During the second cycle, WDE displayed a trending decrease in 2.5-hour food consumption compared to WDU (p = 0.0661), with a significant stress effect [F_(1,58)_ = 4.809, p = 0.0323]. **D-F)** No differences in food intake were detected 24 hours after re-introduction to the WD each week.

### Exposure to a WD and Psychosocial Stress During Adolescence Affects Corticosterone and Testosterone Levels

To clearly delineate the effects of the WD on stress responsivity, we collected fecal samples from each cage to measure concentrations of fecal corticosterone metabolites and fecal testosterone metabolites. Fecal steroid measurements provide a non-invasive method for capturing robust long-term changes (hours to days) in response to stimuli (Sheriff et al., 2010). Fecal samples were collected at four timepoints: **1)** one week after diet introduction, **2)** four weeks after diet introduction, **3)** 24 hours after predator exposure, and **4)** the final day of the study.

There were no differences in fecal corticosterone levels after one week or four weeks on diets **(Figure 5A-B)**. After predator stress exposure, CD animals exhibited elevated corticosterone compared to unexposed controls (CDU:CDE, p = 0.0011; WDU:CDE, p = 0.0001) (**Figure 5C)**. WDE animals showed a significant increase in corticosterone concentration to WDU (p = 0.0260), but not the CDU group (p = 0.2763). Two-way ANOVA revealed significant stress [F_(1,12)_ = 36.57, p < 0.0001] and diet [F_(1,12)_ = 11.24, p = 0.0058] main effects, but no stress x diet interaction effect (p = 0.2122) on corticosterone concentration. Notably, corticosterone concentration was also significantly greater in CDE compared to WDE animals (p = 0.0280). Relative to controls, CDE and WDE animals displayed a 44% and 16% increase in corticosterone release, respectively, in response to the predator stress, implying high-fat diet consumption may blunt corticosterone release in response to stress. On the final week of the study, exposed groups demonstrated heightened fecal corticosterone levels compared to the WDU group (WDU:CDE, p=0.0210; WDU:WDE, p=0.0310) but not the CDU group **(Figure 5D)**.

**Figure 5.**
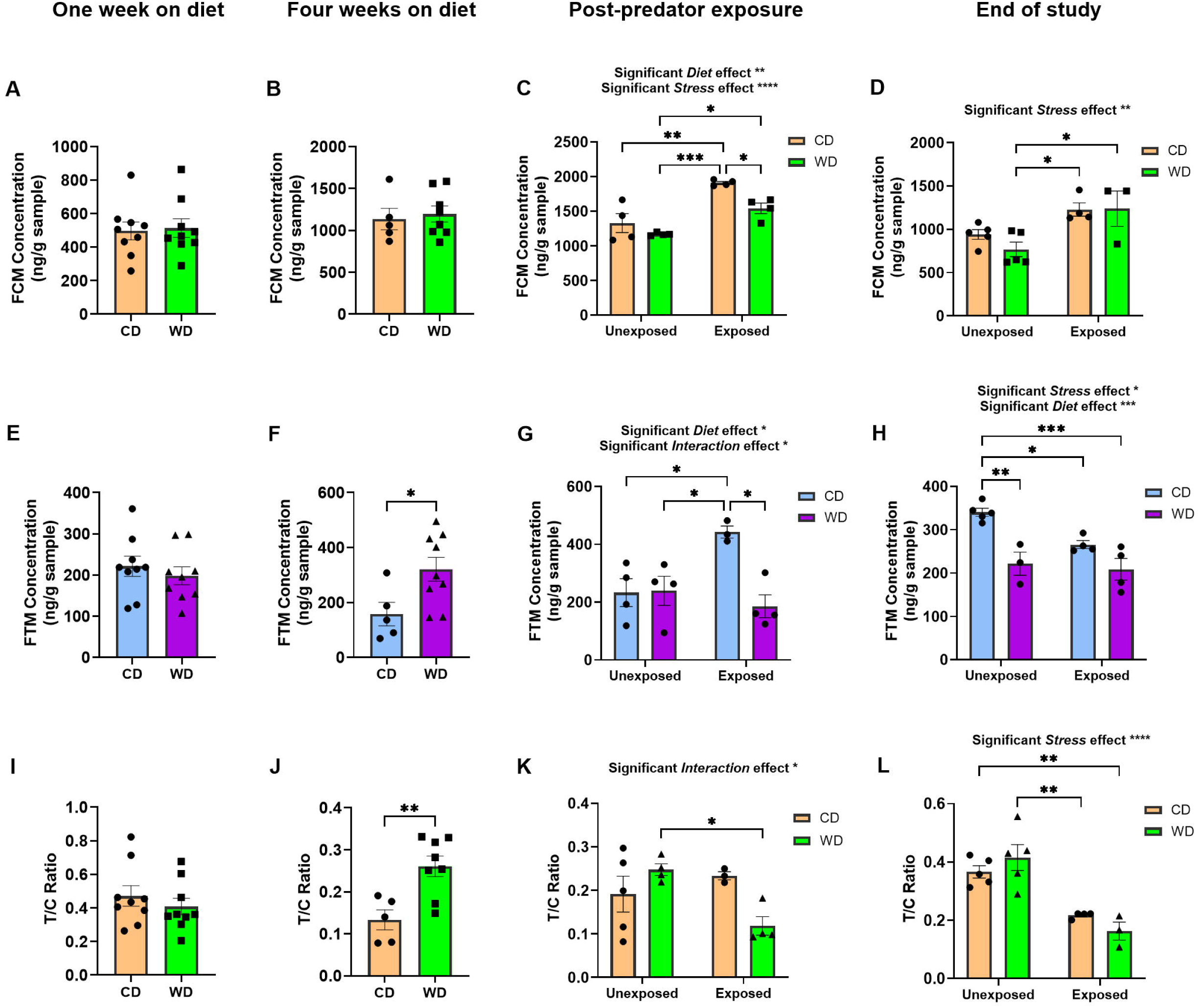
WD and PSS exposure during adolescence dysregulates corticosterone and testosterone levels. **(A-D)** Fecal corticosterone and **(E-H)** testosterone were measured, in ng/g sample, at the following timepoints: 1) One week after diet introduction, 2) four weeks after diet introduction, 3) 24 hours after predator stress, and 4) end of study. **A,E)** There were no differences in early adolescent hormone levels one week after diet introduction or **B)** mid-adolescent corticosterone levels after four weeks on the diets. **F)** Subjects exhibited a significant increase in testosterone levels (p = 0.0312) after four weeks of WD consumption. **C)** Predator-exposed CD rats displayed increased corticosterone compared to unexposed groups (CDE:CDU, p = 0.0011; CDE:WDU, p = 0.0001) and exposed rats that consumed high-saturated-fat diet (CDE:WDE, p = 0.0280) (*stress*, [F_(1,12)_ = 36.57, p < 0.0001]; *diet*, [F_(1,12)_ = 11.24, p = 0.0058]). **G)** Similarly, CDE rats showed an increase in testosterone compared to all other groups, with a significant diet [F_(1,11)_ = 7.917, p = 0.0169] and stress x diet interaction [F_(1,11)_ = 8.766, p = 0.0130] effect. **D)** Young adult rats displayed heightened corticosterone levels following adolescent stress exposure [*stress*, F_(1,13)_ = 14.48, p = 0.0022]. **H)** There was an influence of adolescent stress [F_(1,12)_ = 6.171, p = 0.0287] and diet [F_(1,12)_ = 24.77, p = 0.0003] on decreasing adult testosterone compared to controls. Testosterone-to-corticosterone (T/C) ratio was calculated at each timepoint. **I)** No differences in early adolescent T/C ratio one week after diet introduction. **J)** WD consumption led to an increased T/C ratio (p = 0.0050). **K)** Among rats consuming WD, predator stress exposure decreased T/C ratio (WDE:WDU, p = 0.0413), with a stress main effect [F_(1,12)_ = 8.084, p = 0.0148]. **L)** Adolescent PSS-exposed rats demonstrated a reduced T/C ratio compared to controls [F_(1,13)_ = 39.32, p < 0.0001]. Timepoint 1 and 2: n=5-9 rats/group. Timepoint 3 and 4: n=3-5 rats/group.

There were no differences in testosterone one week after diet introduction **(Figure 5E)**. Four weeks after diet introduction, WD animals showed a significant increase in fecal testosterone metabolite concentration compared to CD animals (p = 0.0312). CD animals subjected to predator stress displayed a significant increase in fecal testosterone concentration compared to all other groups, with a significant diet effect [F_(1,11)_ = 7.917, p = 0.0169] and stress x diet interaction effect [F_(1,11)_ = 8.766, p = 0.0130] **(Figure 5G)**. However, a similar outcome was not observed in exposed rats consuming WD during adolescence. Adolescent high-fat diet and stress exposure resulted in a significant long-term decrease in fecal testosterone metabolites compared to controls, with significant stress [F_(1,12)_ = 6.171, p = 0.0287] and diet [F_(1,12)_ = 24.77, p = 0.0003] main effects, but no stress x diet effect [F_(1,12)_ = 3.075, p = 0.1050] **(Figure 5H)**.

The testosterone-to-corticosterone (T/C) ratio provides an empirical measure for hormonal balance implicated in metabolic balance and physiological state (Romanova et al., 2022; Terburg et al., 2009). This ratio is particularly useful for investigating the effects of chronic stress or resilience factors, as it captures a snapshot of the physiological responses to stress and its capacity for recovery. There was no difference in group T/C ratio after one week on the diet (p = 0.4336) **(Figure 5I)**. WD animals displayed a heightened T/C ratio after four weeks compared to controls (p < 0.0033) **(Figure 5J)**. Two-way ANOVA revealed a stress x diet interaction effect [F_(1,12)_ = 8.084, p = 0.0148] on adolescent T/C ratio following PSS exposure. WDE rats exhibited a decreased T/C ratio compared to WDU animals (p = 0.0413) **(Figure 5K)**. There was a significant main effect of stress on adult T/C ratio [F_(1,13)_ = 39.32, p < 0.0001]. WDE and CDE animals showed a reduced T/C ratio compared to the CDU group (CDU:WDE, p = 0.0041) and WDU group (WDU:CDE, p = 0.0025), respectively **(Figure 5L)**.

### Associations Between Hormonal Levels and Dietary Intake: Impact of Western Diet and Psychosocial Stress on the Endocrine Profiles of Adolescent Rats and its Predictive Validity for Eating Behaviors

We calculated Pearson correlation coefficients to investigate the relationship between corticosterone (C) and testosterone (T) at multiple timepoints across adolescence and young adulthood **(Figure 6)**. Testosterone levels in early and late adolescence showed a moderately strong association with late adolescent corticosterone (C2:T1, r = 0.661, p = 0.014; C2:T2, r = 0.795, p = 0.001). Pre-stress testosterone levels in late adolescence showed a moderate negative association with testosterone in adulthood (T2:T4, r = -0.618, p = 0.032). Predator stress–induced corticosterone levels showed a strong association with adult corticosterone levels (C3:C4, r = 0.704, p = 0.003), validating the robustness of our PSS model to induce long- term endocrine signatures of psychological trauma exposure. After repeating the analyses for both diets separately **(Figure 6B-C)**, we observed a strong inverse relationship between adolescent corticosterone levels and testosterone in late adolescence (C1:T3, r = -0.858, p = 0.014) and young adulthood (C2:T4, r = -0.963, p = 0.008) in CD animals. However, similar associations were not apparent in hormone levels of WD animals.

**Figure 6.**
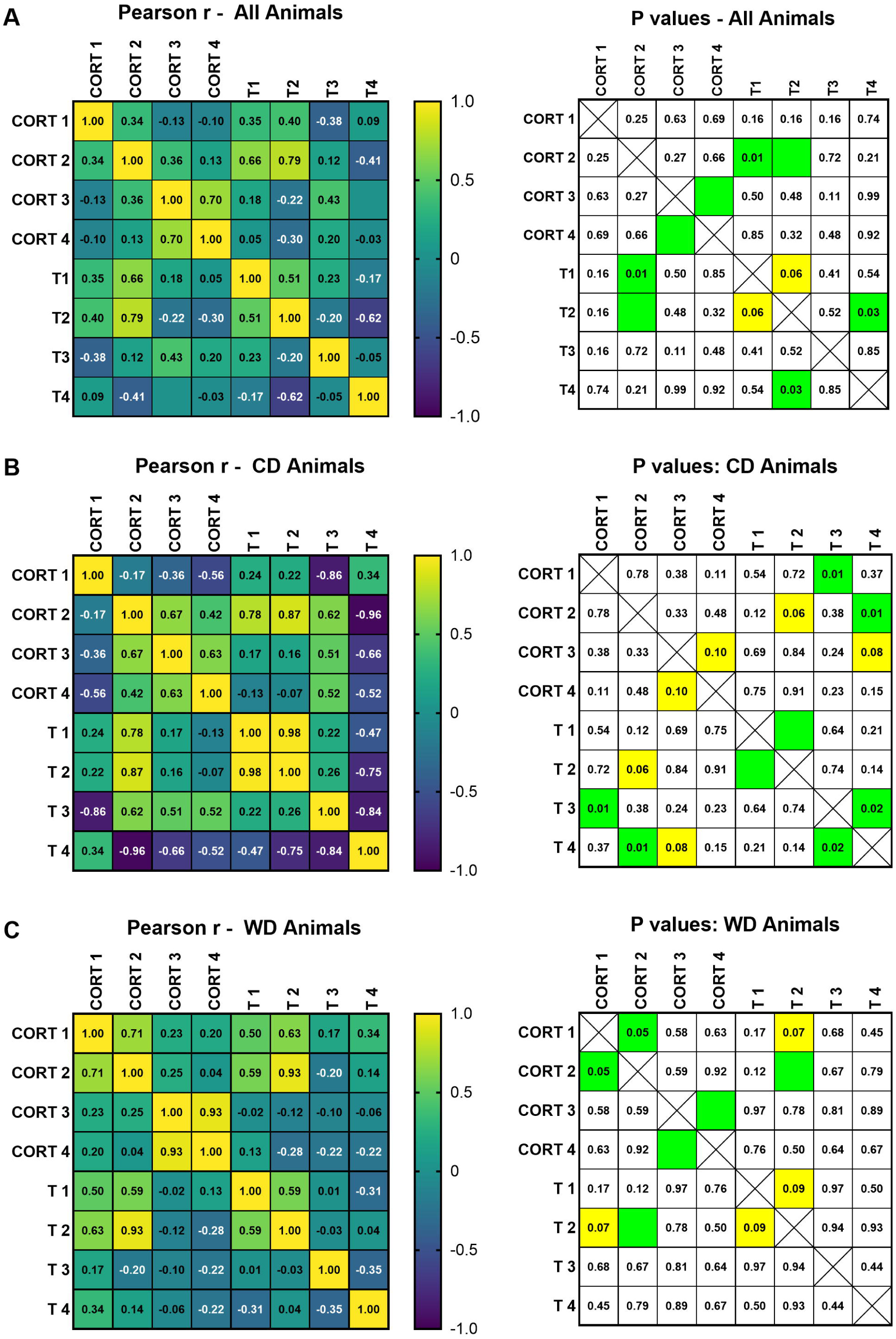
Impact of WD and PSS on the endocrine profiles of adolescent rats. Heatmaps showing Pearson correlation coefficients for corticosterone and testosterone at each timepoint. Correlations were calculated disregarding group **(A)** before repeating the analysis for CD and WD animals separately **(B and C, respectively)**. Corresponding p-values are shown for corresponding correlations displaying significance (p < 0.05, highlighted in green) and trend-like significance (0.05 < p < 0.10, highlighted in yellow). n = 9/

Next, we wanted to determine the association between steroid hormones levels and eating behavior **(S Figure 7-9)**. For CD animals, early adolescent food consumption (FC) was strongly positively associated with adult corticosterone (FC1:C4, r = 0.653, p = 0.057) and late adolescent testosterone (FC1:T3, r = 0.926, p = 0.003) but inversely associated with early adolescent corticosterone (FC1:C1, r = -0.664, p = 0.051) and adult testosterone (FC1:T4, r = - 0.878, p = 0.002) levels. Rats consuming high-fat diet displayed a strong association between early adolescent hormone levels and food consumption in late adolescence (T1:FC2, r = 0.795, p = 0.014) and a trending association in response to acute stress (C1:FC3, r = 0.576, p = 0.104). There was a negative association between early adolescent hormones and bingeing behavior at 2.5-h during BE cycle 1 (C1, r = -0.498, p = 0.035; T1, r = -0.652, p = 0.003).

### Adolescent Hormone Levels Distinguish High and Low Bingeing Phenotypes in Adult Male Rats

Previous investigations from our lab demonstrated that high-saturated fat diet consumption during adolescence increased expression of TACE/ADAM17 (A disintegrin and metalloprotease 17) and IL-6 in brain regions regulating the stress response (Sharafeddin et al., 2024; Vega-Torres et al., 2022). TACE/ADAM17 promotes IL-6 signaling by cleaving the membrane-bound IL-6 receptor (IL-6R), typically only expressed on selective cell types, allowing the receptor to translocate and bind to other cell types (Schumacher and Rose-John, 2019). To further build upon this premise, we performed immunoblotting to measure changes in hippocampal IL-6R. Unexpectedly, no differences were detected in hippocampal IL-6R after four weeks of consuming the high-saturated-fat diet (p = 0.2074) or at the end of the study (p = 0.1840) **(S Figure 10)**. Rats were re-classified into hypophagic or hyperphagic subdivisions based on their 2.5-h food intake during the first binge eating cycle. Independent samples t-test revealed a significant difference in weight-corrected consumption (p < 0.0001) **(S Figure 11)**. We utilized principal component analysis (PCA) to determine whether rats that exhibited increased consumptive behavior expressed similar molecular profiles of early endocrine factors (C1, C2, T1, T2) and adult IL-6R. A score plot of principal component 1 (PC1) vs PC2 was able to separate between hyperphagic and hypophagic rats, with a significant inter-group difference in PC1 (p = 0.0009) but not PC2 (p = 0.2043) scores, indicating that the variables contributing most to PC1 are capable of differentiating between a hyperphagic and hypophagic phenotype **(Figure 7A-C)**. Following this, we wanted to identify the critical factors underlying the variability observed between eating behaviors. We found that IL-6R contributed the least to PC1 and the most to PC2, suggesting that IL-6R does not explain a sufficient portion of the variability in the data. After excluding IL-6R, a re-analysis was performed showing significant differences in PC1 (p = 0.0013) and PC2 (p = 0.0230) between hypophagic and hyperphagic groups **(Figure 7D-F)**. In short, these findings suggest that both early adolescent and mid-late adolescent hormones explain the variation between adult hyperphagic and hypophagic eaters and that adolescent hormone levels may be predictive of susceptibility for disordered eating in adulthood.

**Figure 7.**
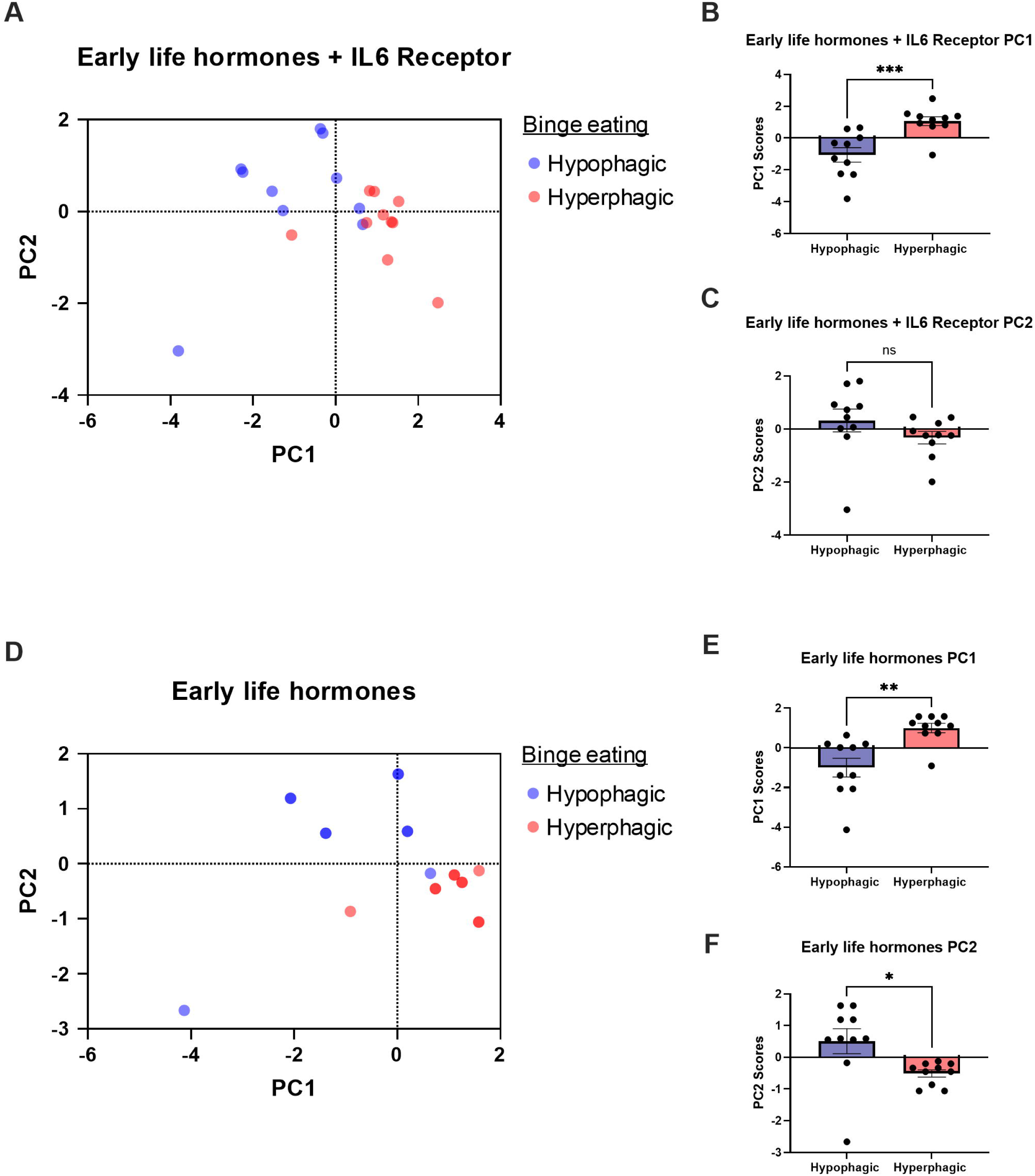
Adolescent hormone levels distinguish between high and low bingeing phenotypes in adult male rats. Subjects were re-classified into subgroups based on 2.5-h consumption during binge eating cycle 1, with the lower half of eaters categorized as hypophagic (blue) and the upper half as hyperphagic (red). For the PCA, corticosterone and testosterone measures at timepoints prior to psychosocial stress exposure (C1, C2, T1, T2) and adult IL-6R expression were included. **A)** A score plot of PC1 and PC2 shows a qualitative separation based on eating behavior. **B,C)** T-test analysis revealed a significant difference in average PC1 scores (p = 0.0009) but not PC2 scores (p = 0.2043) between hypophagic and hyperphagic rats. IL-6R was found to contribute the least to explaining the variability in PC1 but was the highest contributor to PC2. **D)** Therefore, PCA was re-run after removing IL-6R, which strengthened group separation. **E/F)** Both PC1 (p = 0.0013) and PC2 (p = 0.0230) varied significantly between groups, indicating that early hormone levels are associated with binge eating behaviors in adult male rats. n = 10 rats/group.

## DISCUSSION

Obesogenic environments, characterized by access to Western-like high-saturated-fat diets (WD) and psychosocial stress (PSS), are implicated in behavioral and physiological alterations around eating disorders. To our knowledge, the current longitudinal study is the first to evaluate the effects of chronic WD consumption during adolescence on stress resiliency to subsequent development of bingeing behaviors in male Lewis rats. There are four significant findings to report. First, chronic WD consumption led to a rise in adolescent testosterone levels. One of the earliest indicators of pubertal onset is a surge in testosterone due to the activation of the HPG axis. This is significant due to the increasing availability of ‘Westernized’ diets globally and incidents of pubertal development at younger ages. Second, consumption of a WD during adolescence blunts proper endocrine stress responsivity, emphasizing the role of hormone imbalance in modulating stress resiliency. The third significant finding is that adolescent endocrine changes induced by the obesogenic conditions persist into adulthood. This demonstrates the robustness of our model to capture behavioral and metabolic phenotypes. Lastly, our model produces a binge-like phenotype followed by a unique compensation of eating behavior. In particular, rats that endured adolescent obesogenic conditions of high-saturated-fat diet and psychosocial stress displayed hyperphagic tendencies the day following re-introduction to the WD rather than undereat as a consequence of bingeing. Together, these findings validate the impact of traumatic stress on disordered eating behavior. Furthermore, our data supports the role of HPA axis-regulated hormones in mediating diet and stress interactions during the critical period of adolescence.

In the present study, peri-pubertal male Lewis rats (PND50) presented with elevated testosterone after four weeks of access to the WD. This suggests that WD consumption accelerates pubertal timing as initial signs of pubertal development in the Lewis rat typically emerge at PND56 (Flickinger et al., 1997). Prior studies in the lab have demonstrated that WD consumption leads to elevated pro-inflammatory cytokine secretion (Santana et al., 2021; Vega-Torres et al., 2022). The hypothalamus tightly regulates metabolic signaling by employing neuroendocrine factors in response to circulating hormone levels (Cai and Liu, 2011), due to the specialized blood-brain barrier around the ventromedial hypothalamus (Haddad-Tóvolli et al., 2017). However, this makes the hypothalamus uniquely susceptible to overnutrition due to chronic high-fat diet consumption and excess saturated fatty acid intake (Cai and Liu, 2011; Souza et al., 2005; Tzounakou et al., 2024). This is consistent with a growing literature indicating that intake of high saturated fat triggers precocious puberty through hormone imbalance in humans and rodents, although studies are largely conducted with female participants (Calcaterra et al., 2023; Huang et al., 2024; Ullah et al., 2019). Pubertal development begins with stimulation of the HPG axis, initiating a cascade of events eventually resulting in greater gonadal steroid hormone (i.e., testosterone) synthesis and secretion (Sisk and Zehr, 2005). Thus, the balance and timing of steroid hormone fluctuations is crucial for the maturation of the HPA axis and HPG axis. To our knowledge, this is the first longitudinal study evaluating the effects of chronic early high-fat diet consumption on testosterone levels in male Lewis rats.

Our stress paradigm is adapted from a model designed to induce post-traumatic stress disorder–like symptomology using acute (predator odor and live predator exposure) and chronic (housing instability) stressors. Groups exposed to severe acute stress exhibited decreased food intake and greater consumption following chronic social instability stress. Stress can shape eating habits and enhance preference for hyperpalatable food as a coping strategy (Yau and Potenza, 2013). Acute stress induces a physiological response inhibiting appetite (Sominsky and Spencer, 2014), while chronic stress exposure produces a hyperphagic response to palatable foods (Pecoraro et al., 2004). Consistent with the literature, our psychosocial stress model accurately reproduces stress-induced eating phenotypes, providing reliability and relevance of the model to capture concurrent endocrine disruptions.

In this study, groups exposed to severe acute predator stress during late adolescence displayed a substantial increase in corticosterone levels, which was attenuated by high-fat diet consumption. The control diet group exhibited an increase in testosterone, while the high-fat diet group did not produce a significant change in testosterone. Hyperpalatable comfort foods act to reduce activity in the stress response network (Epel et al., 2001; Foster et al., 2009; Gibson, 2006), which may explain the differences in stress responsivity between groups on the control diet or high-fat diet. There is evidence suggesting that stressful experiences alter HPA axis function and that the degree of alteration is dependent on the severity and timing of the stressor (Bosch et al., 2012; Carpenter et al., 2007; Elzinga et al., 2008; McCormick et al., 2010). While glucocorticoid levels are elevated during and immediately after a stressful situation, not much is known about testosterone levels following stress exposure (Romanova et al., 2022). In adults, HPA axis activation typically results in glucocorticoid secretion and subsequent glucocorticoid-mediated suppression of HPG axis activity (Romeo et al., 2005; Stratakis and Chrousos, 1995). Higher testosterone levels are generally associated with lower basal and stress-induced glucocorticoid levels, suggesting a negative association between adrenal and gonadal hormones (Handa et al., 1994; Viau, 2002; Viau and Meaney, 1996). In contrast, testosterone and cortisol levels have been shown to be positively associated during adolescence (Harden et al., 2016). Marceau et al. (2015) proposed that the nascent HPA and HPG axis undergo maturation concurrently during adolescence establishing a positive coupling before assuming a more mature inhibitory relationship. Taken together, this suggests that adolescent consumption of a WD blunts both HPA and HPG axis activity. Further studies are required to delineate specific pathways underlying the deleterious effects of a high-saturated-fat diet on the endocrine stress response.

Intermittent access to a nutritionally complete high-fat diet has been shown to elicit bingeing behavior in rats (Davis et al., 2007). While the binge-eating paradigm did not appear to produce differences in food intake during reintroduction to the high-saturated-fat diet, adolescent exposure to a WD and PSS produced a significant compensatory effect after removal of the diet. This finding is inconsistent with previous binge models using limited access to high-fat diets, in which rats overeat on binge days and undereat on non-binge days (Corwin and Buda-Levin, 2004). Limited access models are capable of producing alterations in body composition and endocrine profile, independent of body weight changes (Blanco-Gandía et al., 2019; Davis et al., 2007). Continuous consumption of an energy-dense diet induces metabolic adaptations, such as increasing lipid metabolism, reducing sensitivity to circulating appetitive hormones, and altering gut microbiota composition, to promote fat storage and weight gain (Roberts et al., 2015; Serino et al., 2012; Sominsky and Spencer, 2014). The current study supports these findings by demonstrating that differences in body weight caused by continuous access to WD are diminished once all groups are placed on a similar diet schedule.

As mentioned previously, groups exposed to psychosocial stress during the adolescent period exhibited elevated corticosterone and reduced testosterone in adulthood weeks after the end of the stress protocol. Altered cortisol reactivity is a measure of vulnerability in patients with stress-related disorders (Girgenti et al., 2017; Zuiden et al., 2013). Introduction to a predator has been demonstrated to impair physiological function and lead to the development of psychopathology in rats after a single exposure (Park et al., 2008). This exemplifies the robustness of our model to produce PTSD-like symptomology. Although chronic adolescent WD consumption led to a similar decrease in testosterone levels as the stress-exposed groups in adulthood, there was no change in corticosterone levels in the absence of stress. This indicates that consumption of a WD may induce long-lasting endocrine changes in the absence of body weight and food consumption differences. Our data demonstrates that the T/C ratio can be relevant to understanding stress-related eating behaviors, weight gain, and metabolic imbalances. Consistent with previous findings (Sharafeddin et al., 2024; Vega-Torres et al., 2022), our study identifies IL-6R as a potential candidate influencing stress-related eating behaviors. Altogether, our results indicate that adolescent high-fat diet consumption and psychosocial stress exposure produce similar long-term endocrine responses, although this appears to be mediated by two separate pathways.

There are several limitations that should be considered for the present study. Firstly, food consumption was assumed to be equally divided among cage partners as animals were housed in pairs. Future studies should incorporate measures of individual metabolic rate to circumvent relying solely on food consumption and to provide more detailed effects of the high-fat diet on eating behavior. Secondly, although this study is among the first to examine the longitudinal effects of diet and stress on hormone fluctuations through adolescence into adulthood, further research is needed to understand how introduction to a high-fat diet at discrete adolescent substages influences diet-induced hormone disruptions and the minimum time required to produce measurable changes in hormone levels due to diet exposure. Understanding the effect of timing of exposure to environmental factors would help identify periods of increased vulnerability within the peri-pubertal phase. Additionally, female rats were excluded from this study limiting the generalizability of our findings. This will be addressed by including female rats in future studies. The inclusion of females would allow us to examine the interaction between sex, puberty, and high-fat diet on stress responses.

Overall, our study highlighted the role of chronic adolescent WD consumption on dysregulated hormone balance in male rats, a phenomenon that is becoming increasingly common as more people adopt a Westernized diet (Drewnowski and Popkin, 1997). Obesogenic environments, with easy access to palatable food and daily stressors, are becoming more pervasive. These environments can initiate the stress response and perpetuate it for prolonged periods of time. However, deficits in HPA axis related brain structures may not be apparent until adulthood (Isgor et al., 2004). This underscores the need to develop early predictive biomarkers for more vulnerable individuals to prevent and reduce risk of psychopathology. The deficits in stress responsivity elicited by chronic WD intake reflect human conditions of heightened susceptibility to psychopathology during an increasingly sensitive period of development. Further studies are required to elucidate the mechanisms involved in the interplay of diet and stress and the role of hormones as mediators between diet and stress effects during adolescence and the long-term outcomes that persist into adulthood.

### Data availability statement

In addition to the data presented in the supplementary materials, supportive datasets are available from the corresponding author upon reasonable request.

## Acknowledgements

We would like to acknowledge the funding provided for this research project from the NIH (DK124727, GM060507, and MD006988) and the Loma Linda University School of Medicine GRASP Seed Funds awarded to JDF. We would like to thank the Loma Linda University Animal Care Facility staff and the cat owner who generously allowed their pet to participate in this study. Thank you to the Center for Health Disparities and Molecular Medicine staff. A special thanks to Vivianna Williams and Giara Wright for research assistance. We would also like to extend a special acknowledgement to Jasmine the cat, whose participation was instrumental in generating the stress model.

## CRediT authorship contribution statement

**Julio Sierra:** Conceptualization, Formal analysis, Investigation, Methodology, Validation, Visualization, Writing – original draft preparation, Writing – review and editing. **Timothy B. Simon:** Conceptualization, Formal analysis, Investigation, Methodology, Supervision, Validation, Visualization, Writing – review and editing. **Darine Abu Hilal**: Investigation, Visualization, Writing – review and editing. **Yaria Arroyo Torres:** Formal analysis, Investigation, Methodology. **Jose M. Santiago Santana:** Formal analysis, Investigation, Methodology, Resources. **Johnny D. Figueroa:** Conceptualization, Formal analysis, Funding acquisition, Methodology, Project administration, Resources, Supervision, Validation, Writing – review and editing.

