## Supplemental Materials for "Impact of adolescent high-fat diet and psychosocial stress on neuroendocrine stress responses and binge eating behavior in adult male Lewis rats"

**Supplemental Figure 1. Longitudinal weight-corrected food consumption.** Weekly weight-corrected food consumption for control diet (CD) and Western diet (WD) groups (Week 1-4). Groups were subdivided into PSS-exposed (E) and unexposed (U) subgroups after four weeks. Rats were subjected to psychosocial stress for 11 days (pink bar). Differences in food consumption were observed every week, except the week after exposure to a severe predator stressor (week 5). Week 1-4: sample size n=38-40 rats/group, Week 5-10: n=16 rats/group * p<0.05, ** p<0.01, *** p<0.001.

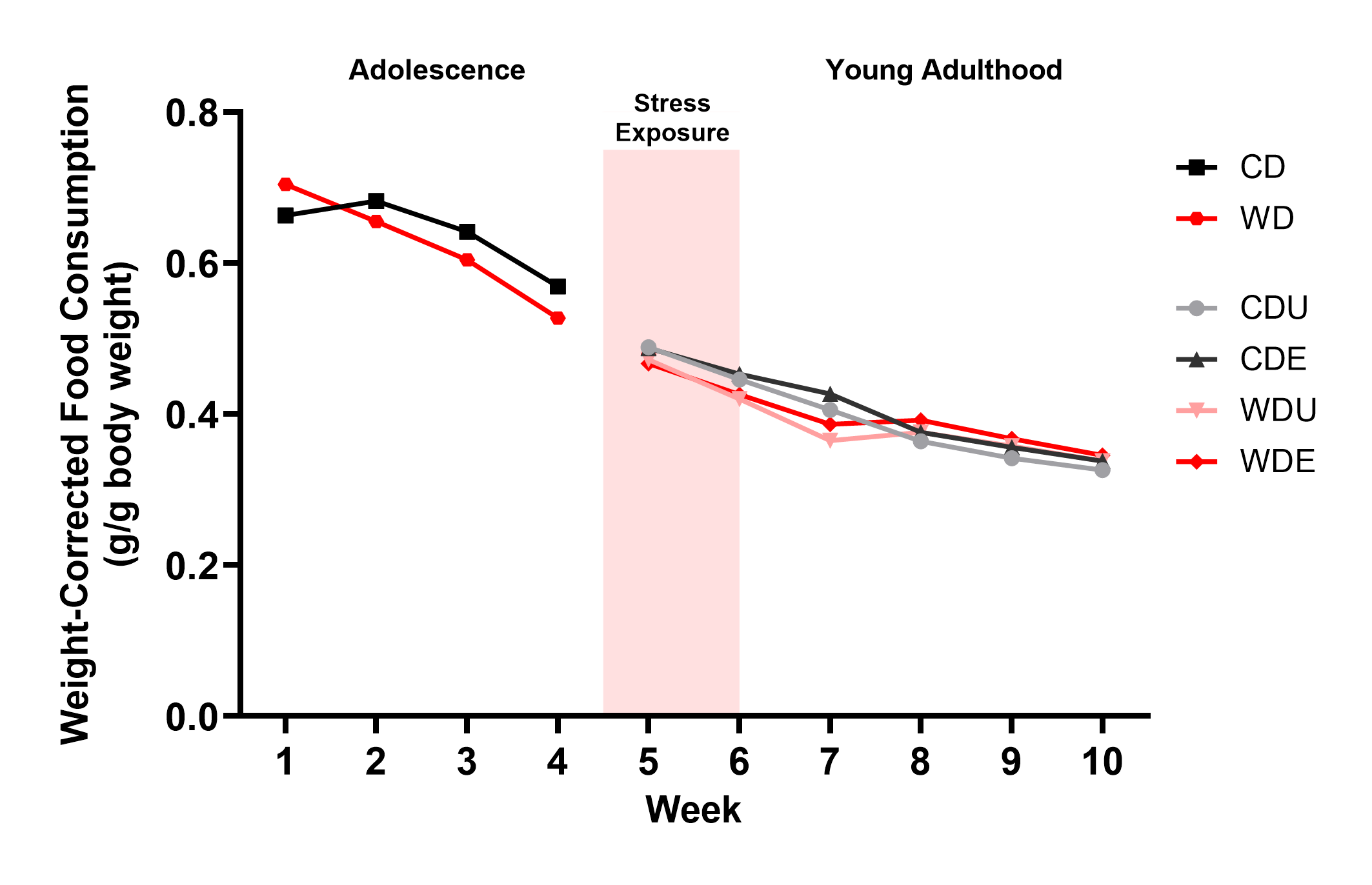

**Supplemental Figure 2. Changes in food intake following exposure to a live predator. A)** Adolescent rats that consumed Western-like high-saturated-fat diet (WD) and were exposed to a severe acute predator stressor exhibited decreased food intake compared to unexposed WD rats (p = 0.0030). There was a significant main effect of stress on food intake [F_(1,60)_ = 13.36, p = 0.0005]. **B)** After correcting for body weight differences, we observed a significant effect of diet on weight-corrected food intake [F_(1,59)_ = 5.585, p = 0.0214]. *Post hoc* analysis did not reveal differences in corrected-food intake between groups.

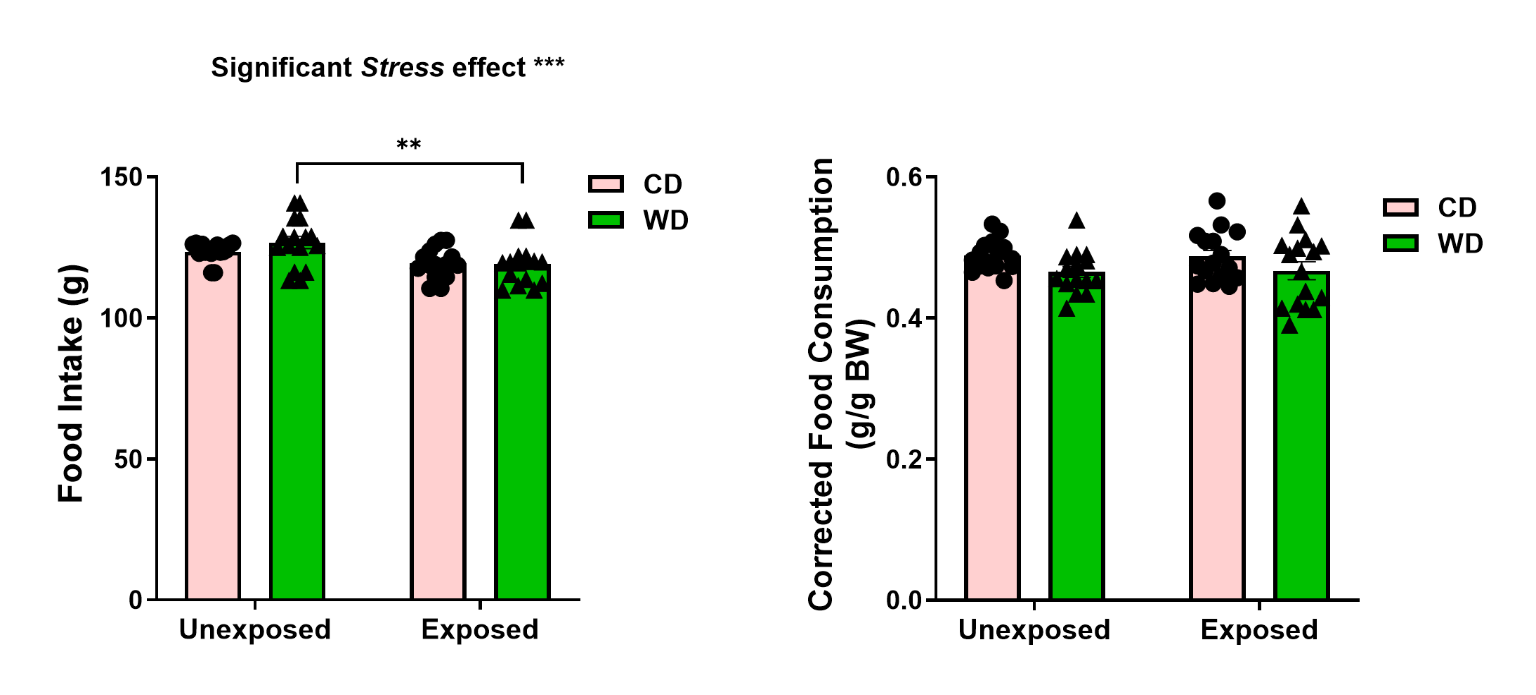

**Supplemental Figure 3. Food intake following social instability stress.** Weight-corrected food intake was affected by diet [F_(1,60)_ = 17.54, p < 0.0001] but not stress [F_(1,60)_ = 1.038, p = 0.3124] following social instability, with WD animals showing decreased food consumption compared to CD animals. CDE animals ate significantly more than WDU animals (p = 0.0030).

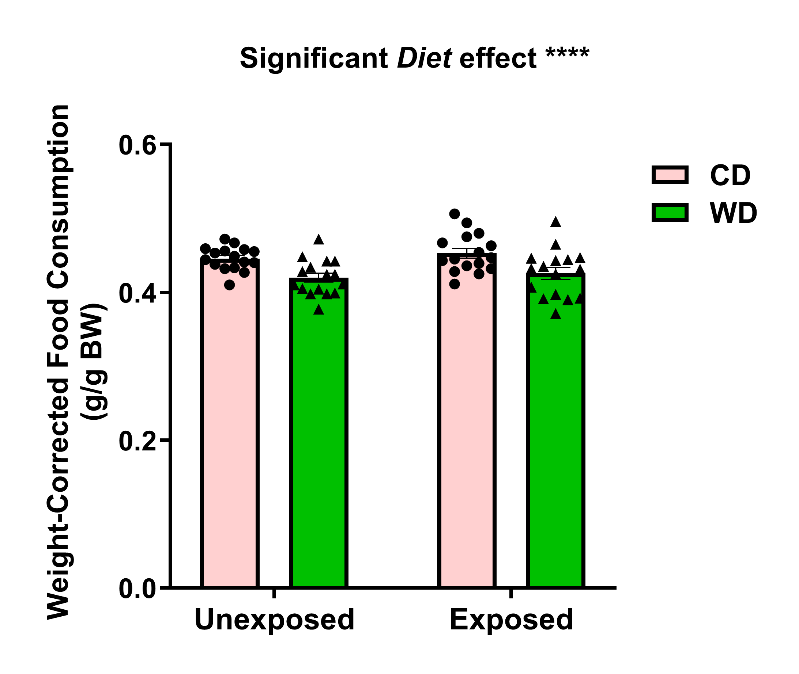

**Supplemental Figure 4. Impact of psychosocial stress (PSS) on food intake.** Across the entire PSS protocol, there was a significant main effect of diet [F_(1,60)_ = 11.12, p = 0.0015]. Group analysis showed significantly greater consumption in the CDE group compared to the WDE group (p = 0.0351).

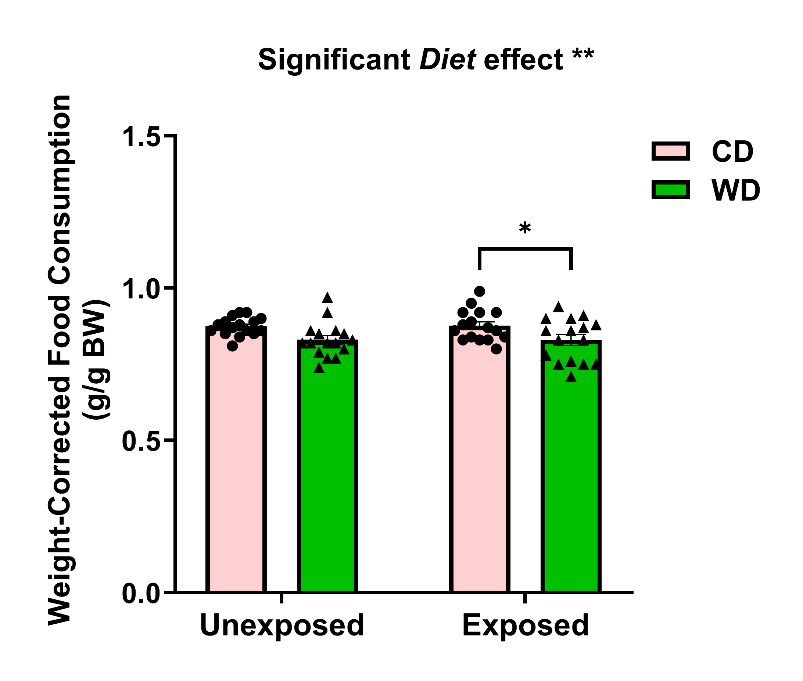

**Supplemental Figure 5. 24-h consumption following re-introduction to CD during binge eating cycles.** We observed a consistent effect of stress on 24-hour food consumption after replacing WD with CD throughout the binge eating cycles (*cycle 1*, [F_(1,60)_ = 3.793, p = 0.0562]; *cycle 2*, [F_(1,60)_ = 3.380, p = 0.0709]; *cycle 3*, [F_(1,60)_ = 9.795, p = 0.0027]). Further, the WDE group shows significantly increased consumption compared to controls during the first (p = 0.0061) and third cycles (p = 0.0157), with a trend towards significance in the second cycle (p = 0.0599).

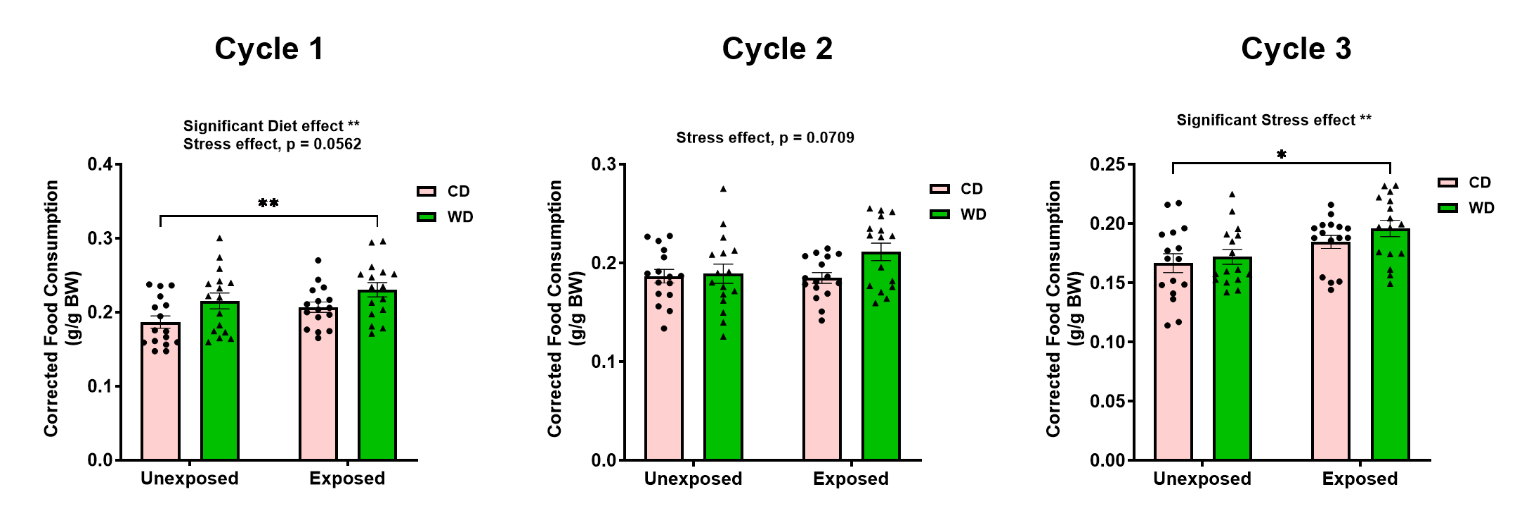

**Supplemental Figure 6. Weekly food consumption during binge eating cycles.** Weekly food consumption during the binge eating paradigm was influenced by diet during all three cycles (*cycle 1*, [F_(1,60)_ = 6.991, p = 0.0095]; *cycle 2*, [F_(1,60)_ = 8.931, p = 0.0041]; *cycle 3*, [F_(1,60)_ = 4.299, p = 0.0425]). Additionally, there was a significant effect of stress on food consumption during cycle 1 [F_(1,60)_ = 6.991, p = 0.0104]. WDE rats displayed increased consumption compared to CDU rats during the first (p = 0.0021) and second (p = 0.0114) binge eating cycles.

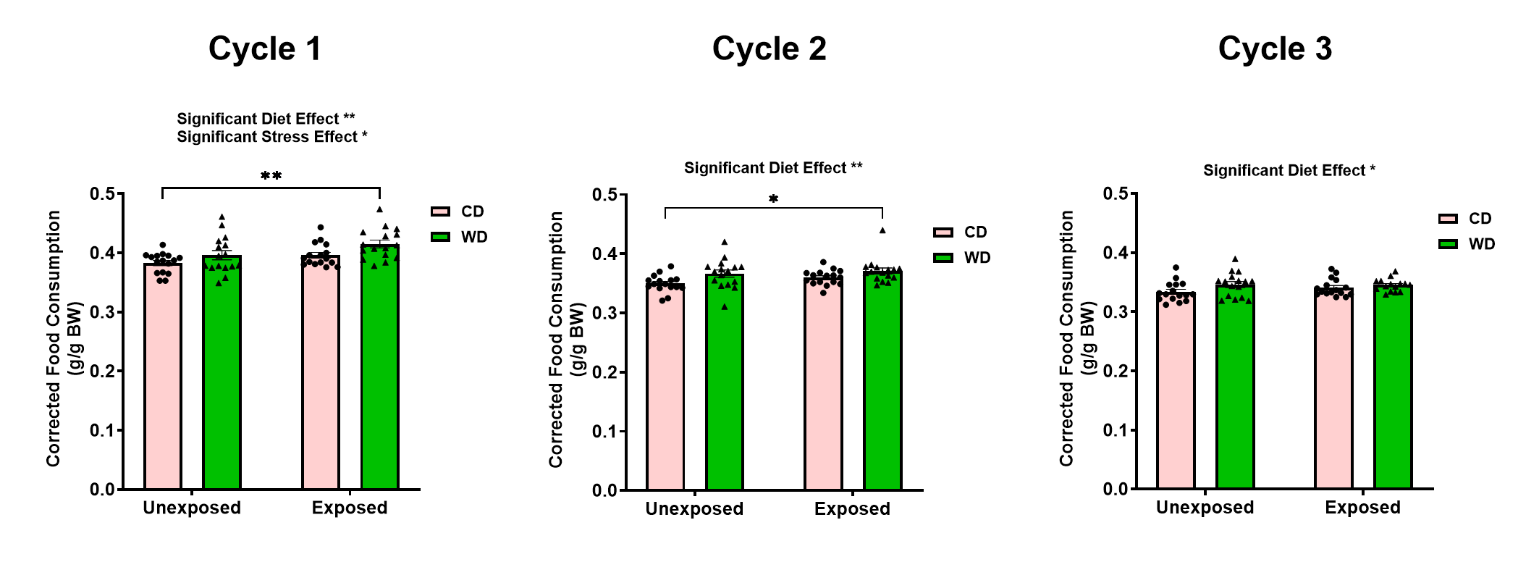

**Supplemental Figure 7. Association between corticosterone, testosterone, and corrected food consumption in male rats that consumed a low-fat control diet (CD).** Pearson correlation analysis shows the relationship between corticosterone, testosterone, and weekly food consumption taken from the same week samples were collected for CD animals, regardless of stress exposure. Sample size n = 7-9 after removing outliers.

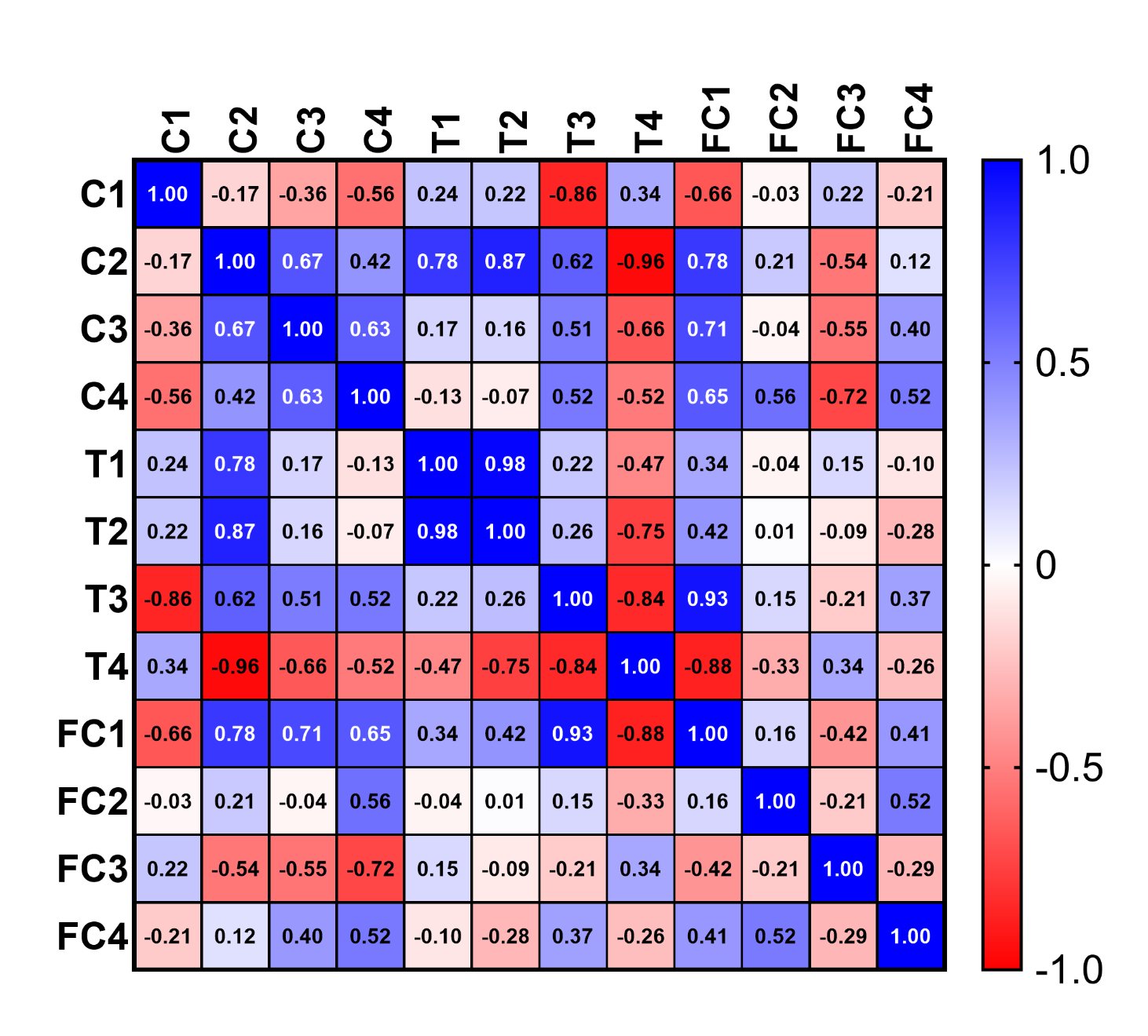

**Supplemental Figure 8. Association between corticosterone, testosterone, and corrected food consumption in male rats that consumed a Western-like high-saturated-fat diet (WD).** Pearson correlation analysis shows the relationship between corticosterone, testosterone, and weekly food consumption taken at the corresponding timepoints for WD animals, regardless of stress exposure. Sample size n = 7-9 after removing outliers.
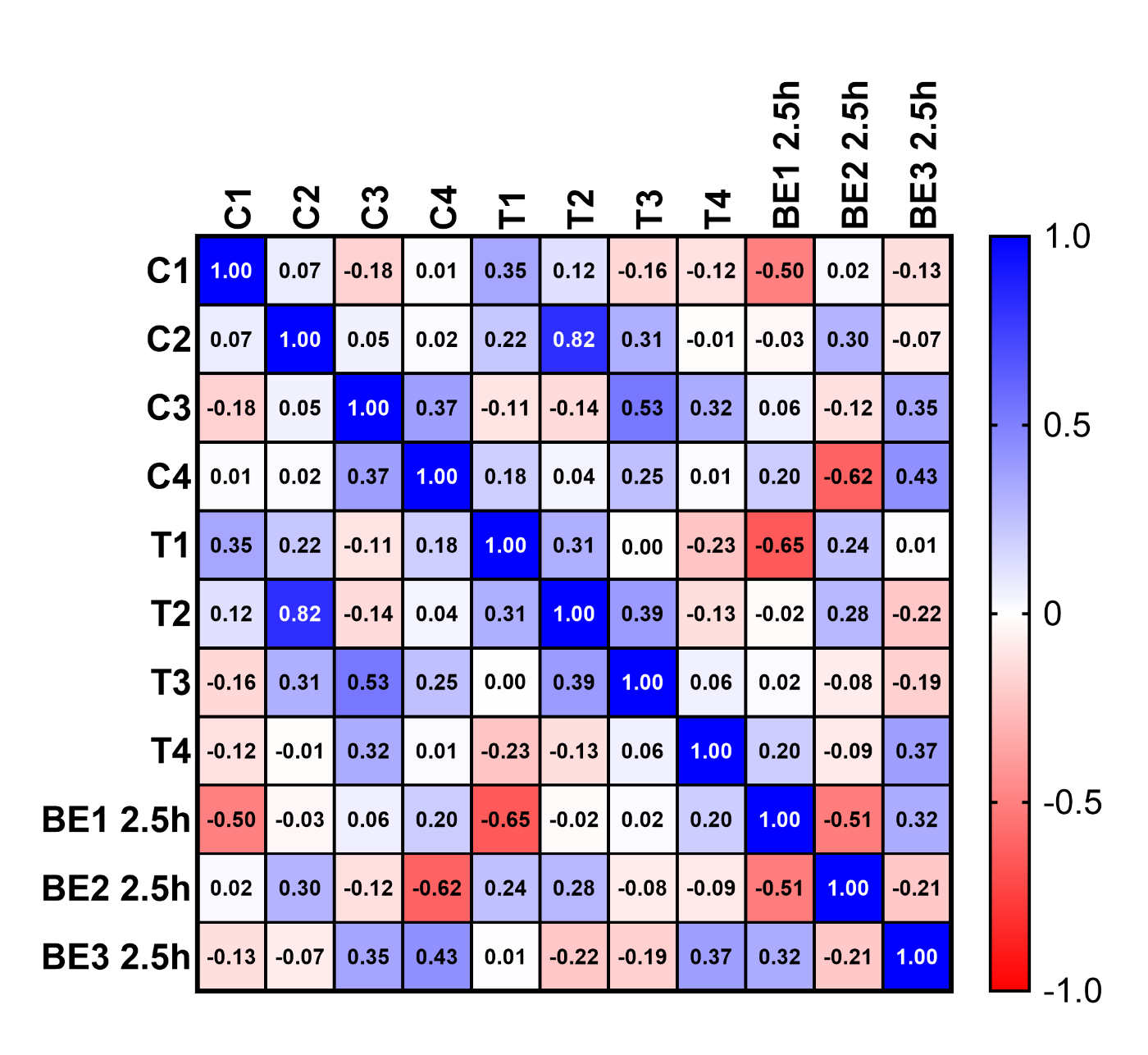

**Supplemental Figure 9. Correlation matrix of hormones and food consumption during binge eating cycles.** Pearson correlation analysis shows the relationship between corticosterone, testosterone, and binging behavior. Binging behavior is defined by 2.5-h WD consumption after (re-)introduction to the WD during the binge eating paradigm. Sample size n = 18.

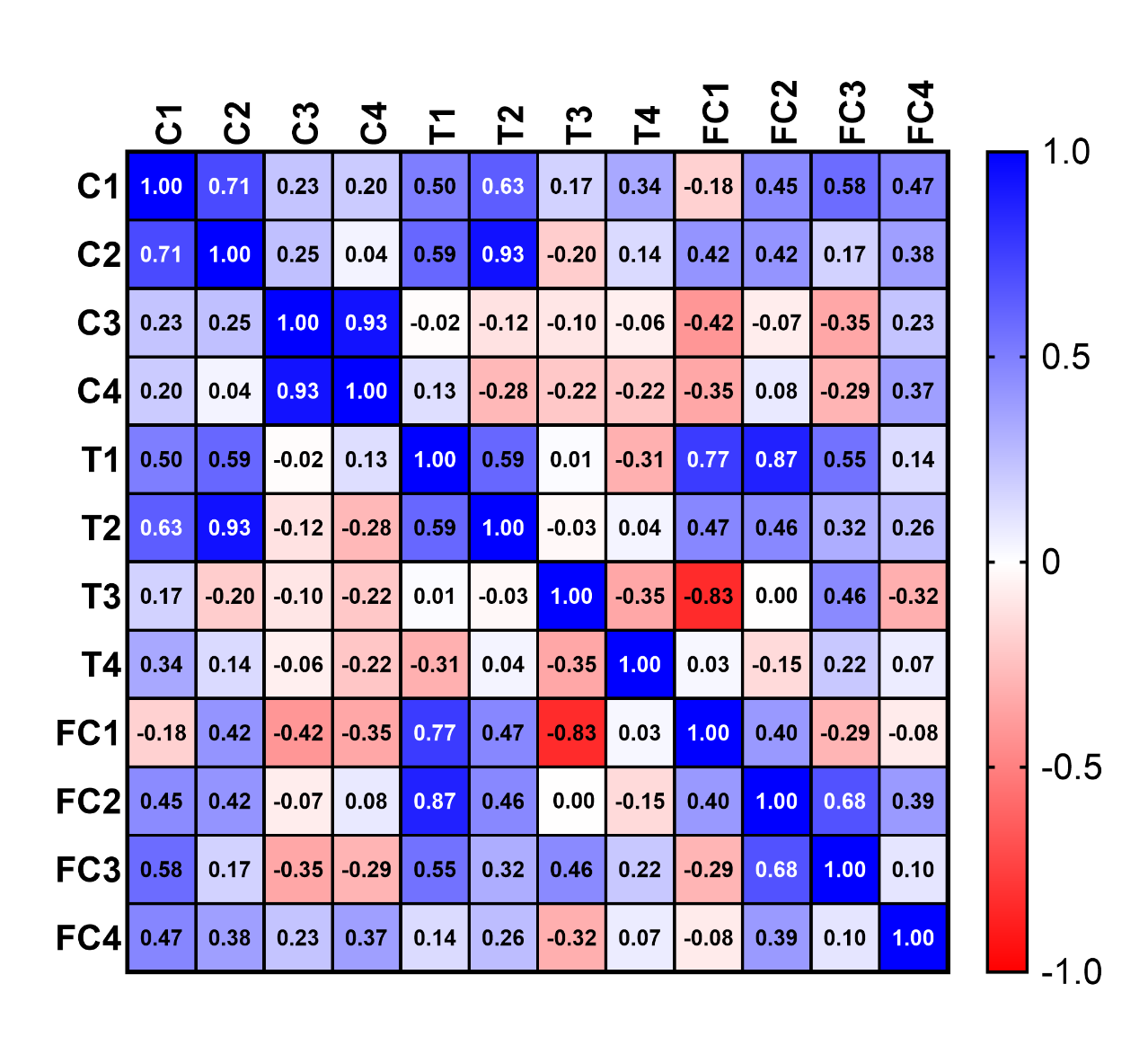

**Supplemental Figure 10. Changes in IL-6R expression following diet and psychosocial stress exposure.** IL-6R (80 kDa) expression was measured by Western immunoblot in a subset of animals **A)** four weeks after diet introduction and **B)** at the end of the study. No group differences were observed between diet groups or the treatment of diet and stress. Sample size n = 4-5 rats/group.

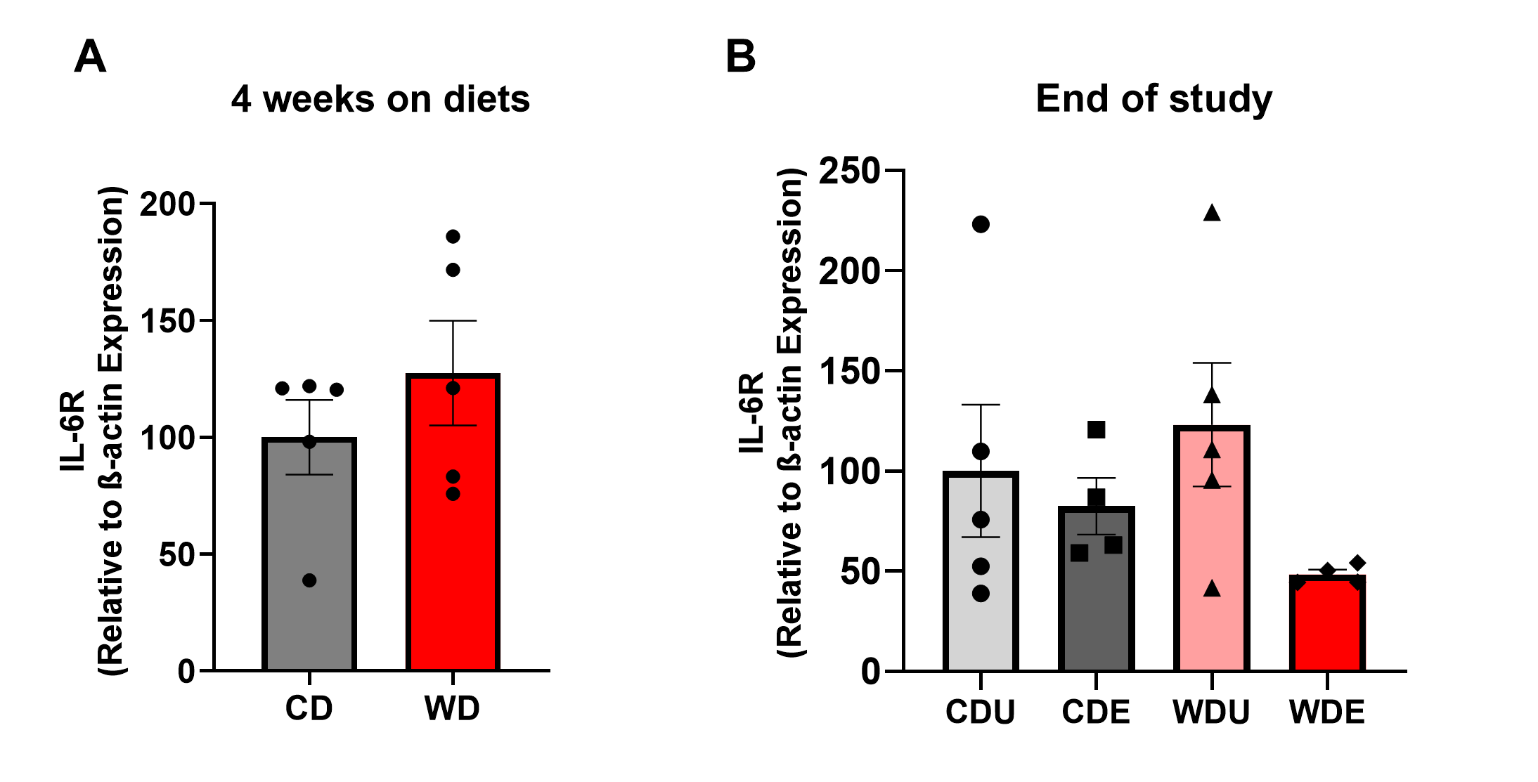

**Supplemental Figure 11. Differences in binging behavior during cycle 1 of the binge eating paradigm.** Animals were re-classified into hypophagic and hyperphagic groups, regardless of diet or stress exposure, using 2.5-h consumption during binge eating cycle 1. Hyperphagic animals displayed significantly higher consumption than hypophagic animals (p < 0.0001). Sample size n = 10 rats/group.

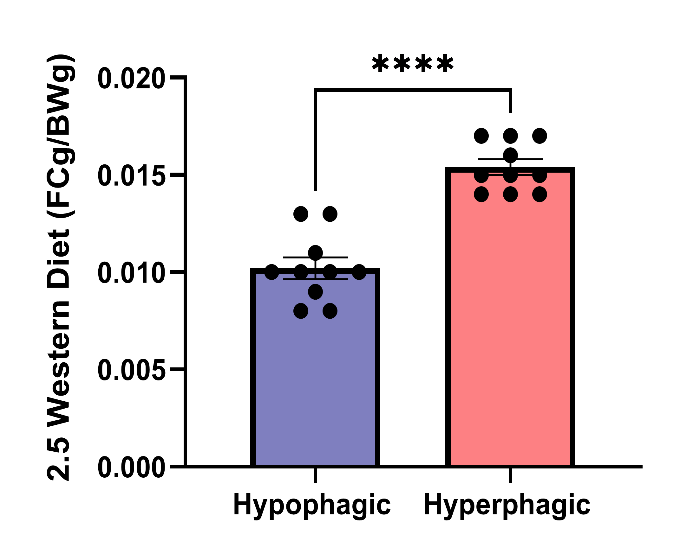

**Supplemental Table 1. Macronutrient composition of the custom purified diets.** Detailed composition of macronutrients for the ingredient-matched low-fat purified control diet (CD) and Western-like high-saturated-fat diet (WD). Below is a detailed description of the fatty acids (in g/kg) per diet.

| **Macronutrient** | **CD** | **WD** |
| --- | --- | --- |
| % kcal Carbohydrates (corn starch) | 64.7 | 43.1 |
| % kcal Protein (casein) | 18.8 | 15.5 |
| % kcal Fat (milk fat) | 16.5 | 41.4 |
| % kcal Total kcal | 3.8 | 4.6 |
| **Fatty Acid Class** | **g/kg Diet** | **g/kg Diet** |
| Butryic (C4:0) | 0.70 | 4.43 |
| Caproic (6:0) | 0.53 | 3.28 |
| Caprylic (C8:0) | 0.34 | 2.12 |
| Capric (C10:0) | 0.81 | 5.19 |
| Lauric (C12:0) | 0.10 | 6.29 |
| Myristic (C14:0) | 3.09 | 19.5 |
| Myristoleic (C14:1) | 0.27 | 1.70 |
| Pentadecanoic (15:0) | 0.34 | 2.12 |
| Palmitic (C16:0) | 11.8 | 59.4 |
| Palmitoleic (C16:1) | 0.47 | 2.91 |
| Heptadecanoic (17:0) | 0.17 | 1.02 |
| Stearic (C18:0) | 3.11 | 17.4 |
| Oleic (C18:1) | 10.5 | 34.9 |
| Linoleic (C18:2) | 11.4 | 9.73 |
| Alpha Linolenic (C18:3) | 0.33 | 0.92 |
| Arachidic (20:0) | 0.11 | 0.25 |
| Homogamma Linolenic (20:3) | <0.07 | 0.19 |
| Arachidonic (n6) 20:4 | <0.07 | 0.26 |
| EPA (20:5 n-3) | <0.07 | <0.07 |
| DHA (22:6 n-3) | <0.07 | <0.07 |
| SFA | 20.9 | 114 |
| MUFA | 11.8 | 44.0 |
| PUFA | 11.3 | 10.8 |
| Omega 3 Fatty Acids | 0.33 | 1.11 |
| Omega 6 Fatty Acids | 11.4 | 10.2 |

**Supplemental Table 2.** Weekly body weight statistics. Body weight measurements were taken weekly. Two-way repeated-measures ANOVA was performed, with diet and time as factors, during the first four weeks. A TW-RM ANOVA was used to evaluate the data from week 5 after groups underwent exposure to psychosocial stress to the end of the study, with time and treatment (diet + stress) as factors.

| **ANOVA Table** | **SS** | **DF** | **MS** | **F (DFn, DFd)** | **P Value** |
| --- | --- | --- | --- | --- | --- |
| **Time x Diet** | **1720** | **4** | **430.1** | **F (4, 304) = 20.19** | **P<0.0001** |
| **Time** | **1482523** | **4** | **370631** | **F (1.353, 102.8) = 17396** | **P<0.0001** |
| Diet | 3861 | 1 | 3861 | F (1, 76) = 6.884 | P=0.0105 |
| **Subject** | **42627** | **76** | **560.9** | **F (76, 304) = 26.33** | **P<0.0001** |
| Residual | 6477 | 304 | 21.3 |  |  |

| **Test Details** | **Mean 1** | **Mean 2** | **Mean Diff.** | **SE of Diff** | **N1** | **N2** | **t** | **DF** | **95.00% Cl of diff** | **Adjusted p-value** |
| --- | --- | --- | --- | --- | --- | --- | --- | --- | --- | --- |
| CD - WD |  |  |  |  |  |  |  |  |  |  |
| Week 0 | 42.77 | 42.66 | 0.1187 | 1.128 | 38 | 40 | 0.1052 | 75.93 | -2.854 to 3.091 | >0.9999 |
| Week 1 | 73.13 | 77.46 | -4.329 | 2.048 | 38 | 40 | 2.113 | 74.94 | -9.727 to 1.070 | 0.1758 |
| Week 2 | 117.5 | 123.3 | -5.847 | 2.683 | 38 | 40 | 2.179 | 75.99 | -12.92 to 1.222 | 0.152 |
| **Week 3** | 162.2 | 171.5 | -9.274 | 2.954 | 38 | 40 | 3.139 | 75.79 | **-17.06 to -1.490** | **0.012** |
| **Week 4** | 208.4 | 220.6 | -12.14 | 3.407 | 38 | 40 | 3.564 | 75.27 | **-21.12 to -3.165** | **0.0032** |

| **ANOVA Table** | **SS** | **DF** | **MS** | **F (DFn, DFd)** | **P Value** |
| --- | --- | --- | --- | --- | --- |
| Time x Group | 1155 | 15 | 77 | F (15, 300) = 2.761 | P=0.0005 |
| Time | 805011 | 5 | 161002 | F (2.185, 131.1) = 5772 | P<0.0001 |
| Group | 28284 | 3 | 9428 | F (3, 60) = 3.797 | P=0.0147 |
| Subject | 148991 | 60 | 2483 | F (60, 300) = 89.03 | P<0.0001 |
| Residual | 8368 | 300 | 27.89 |  |  |

| **Test Details** | **Mean 1** | **Mean 2** | **Mean**  **Diff.** | **SE of Diff** | **N1** | **N2** | **q** | **DF** | **95% Cl of diff** | **Adjusted  p-value** |
| --- | --- | --- | --- | --- | --- | --- | --- | --- | --- | --- |
| Week 5 |  |  |  |  |  |  |  |  |  |  |
| CDU vs. CDE | 253.3 | 245.9 | 7.400 | 5.500 | 16 | 16 | 1.903 | 24.79 | -7.738 to 22.54 | 0.5439 |
| **CDU vs. WDU** | 253.3 | 269.2 | -15.86 | 4.997 | 16 | 16 | 4.489 | 26.83 | **-29.54 to -2.181** | **0.0185** |
| CDU vs. WDE | 253.3 | 256.6 | -3.300 | 5.648 | 16 | 16 | 0.8263 | 24.26 | -18.87 to 12.27 | 0.9359 |
| **CDE vs. WDU** | 245.9 | 269.2 | -23.26 | 6.232 | 16 | 16 | 5.279 | 29.46 | **-40.23 to -6.298** | **0.0043** |
| CDE vs. WDE | 245.9 | 256.6 | -10.70 | 6.765 | 16 | 16 | 2.237 | 29.96 | -29.10 to 7.697 | 0.404 |
| WDU vs. WDE | 269.2 | 256.6 | 12.56 | 6.363 | 16 | 16 | 2.792 | 29.15 | -4.769 to 29.89 | 0.2208 |
| Week 6 |  |  |  |  |  |  |  |  |  |  |
| CDU vs. CDE | 287.9 | 281.9 | 5.950 | 5.671 | 16 | 16 | 1.484 | 24.76 | -9.660 to 21.56 | 0.7227 |
| **CDU vs. WDU** | 287.9 | 309.7 | -21.85 | 5.747 | 16 | 16 | 5.377 | 24.49 | **-37.68 to -6.019** | **0.0044** |
| CDU vs. WDE | 287.9 | 295.8 | -7.875 | 6.328 | 16 | 16 | 1.760 | 22.72 | -25.40 to 9.654 | 0.6061 |
| **CDE vs. WDU** | 281.9 | 309.7 | -27.80 | 6.916 | 16 | 16 | 5.685 | 29.99 | **-46.60 to -8.995** | **0.0019** |
| CDE vs. WDE | 281.9 | 295.8 | -13.83 | 7.406 | 16 | 16 | 2.640 | 29.39 | -33.99 to 6.335 | 0.2639 |
| WDU vs. WDE | 309.7 | 295.8 | 13.98 | 7.464 | 16 | 16 | 2.648 | 29.53 | -6.339 to 34.29 | 0.2614 |
| Week 7 |  |  |  |  |  |  |  |  |  |  |
| CDU vs. CDE | 318.2 | 314.1 | 4.125 | 5.839 | 16 | 16 | 0.9990 | 26.89 | -11.86 to 20.11 | 0.8937 |
| CDU vs. WDU | 318.2 | 333.4 | -15.23 | 6.407 | 16 | 16 | 3.36 | 24.91 | -32.85 to 2.403 | 0.1081 |
| CDU vs. WDE | 318.2 | 324.6 | -6.412 | 6.893 | 16 | 16 | 1.316 | 23.49 | -25.46 to 12.63 | 0.7890 |
| CDE vs. WDU | 314.1 | 333.4 | -19.35 | 7.256 | 16 | 16 | 3.771 | 29.49 | -39.10 to 0.4007 | 0.0565 |
| CDE vs. WDE | 314.1 | 324.6 | -10.54 | 7.689 | 16 | 16 | 1.938 | 28.53 | -31.51 to 10.43 | 0.5274 |
| WDU vs. WDE | 333.4 | 324.6 | 8.813 | 8.129 | 16 | 16 | 1.533 | 29.72 | -13.30 to 30.93 | 0.7017 |
| Week 8 |  |  |  |  |  |  |  |  |  |  |
| CDU vs. CDE | 342.5 | 339.3 | 3.213 | 6.583 | 16 | 16 | 0.6902 | 26.97 | -14.80 to 21.23 | 0.9611 |
| CDU vs. WDU | 342.5 | 359.5 | -16.98 | 6.86 | 16 | 16 | 3.499 | 26.08 | -35.79 to 1.841 | 0.0879 |
| CDU vs. WDE | 342.5 | 352.7 | -10.19 | 7.330 | 16 | 16 | 1.966 | 24.69 | -30.37 to 9.991 | 0.5172 |
| CDE vs. WDU | 339.3 | 359.5 | -20.19 | 7.847 | 16 | 16 | 3.638 | 29.89 | -41.53 to 1.155 | 0.069 |
| CDE vs. WDE | 339.3 | 352.7 | -13.40 | 8.261 | 16 | 16 | 2.294 | 29.32 | -35.89 to 9.092 | 0.3823 |
| WDU vs. WDE | 359.5 | 352.7 | 6.787 | 8.484 | 16 | 16 | 1.131 | 29.75 | -16.29 to 29.87 | 0.8538 |
| Week 9 |  |  |  |  |  |  |  |  |  |  |
| CDU vs. CDE | 363.3 | 362.1 | 1.175 | 7.059 | 16 | 16 | 0.2354 | 28.58 | -18.07 to 20.42 | 0.9983 |
| CDU vs. WDU | 363.3 | 382.2 | -18.91 | 7.77 | 16 | 16 | 3.442 | 26.58 | -40.20 to 2.372 | 0.0949 |
| CDU vs. WDE | 363.3 | 376.6 | -13.31 | 8.076 | 16 | 16 | 2.331 | 25.75 | -35.48 to 8.857 | 0.3706 |
| CDE vs. WDU | 362.1 | 382.2 | -20.09 | 8.454 | 16 | 16 | 3.36 | 29.36 | -43.10 to 2.930 | 0.1042 |
| CDE vs. WDE | 362.1 | 376.6 | -14.49 | 8.736 | 16 | 16 | 2.345 | 28.83 | -38.30 to 9.324 | 0.3635 |
| WDU vs. WDE | 382.2 | 376.6 | 5.6 | 9.32 | 16 | 16 | 0.8497 | 29.91 | -19.75 to 30.95 | 0.931 |
| Week 10 |  |  |  |  |  |  |  |  |  |  |
| CDU vs. CDE | 382.8 | 381.4 | 1.400 | 7.236 | 16 | 16 | 0.2736 | 28.55 | -18.33 to 21.13 | 0.9974 |
| CDU vs. WDU | 382.8 | 403.6 | -20.79 | 8.043 | 16 | 16 | 3.655 | 26.34 | -42.83 to 1.258 | 0.0698 |
| CDU vs. WDE | 382.8 | 397.3 | -14.43 | 8.27 | 16 | 16 | 2.467 | 25.74 | -37.13 to 8.278 | 0.3225 |
| CDE vs. WDU | 381.4 | 403.6 | -22.19 | 8.745 | 16 | 16 | 3.588 | 29.24 | -46.00 to 1.626 | 0.0748 |
| CDE vs. WDE | 381.4 | 397.3 | -15.83 | 8.954 | 16 | 16 | 2.499 | 28.85 | -40.23 to 8.579 | 0.3092 |
| WDU vs. WDE | 403.600 | 397.3 | 6.363 | 9.618 | 16 | 16 | 0.9355 | 29.95 | -19.79 to 32.52 | 0.9106 |

**Supplemental Table 3.** Weekly weight-corrected food consumption. Food consumption was measured weekly. Two-way repeated-measures ANOVA was performed, with diet and time as factors, during the first four weeks. A TW-RM ANOVA was used to evaluate the data from week 5 after groups underwent exposure to psychosocial stress to the end of the study, with time and treatment (diet + stress) as factors.

| **ANOVA Table** | **SS** | **DF** | **MS** | **F (DFn, DFd)** | **Adjusted p-value** |
| --- | --- | --- | --- | --- | --- |
| **Time x Diet** | **0.08818** | **3** | **0.02939** | **F (3, 228) = 51.82** | **P<0.0001** |
| **Time** | **0.8668** | **3** | **0.2889** | **F (2.029, 154.2) = 509.4** | **P<0.0001** |
| Diet | 0.02053 | 1 | 0.02053 | F (1, 76) = 3.760 | P=0.0562 |
| **Subject** | **0.415** | **76** | **0.00546** | **F (76, 228) = 9.627** | **P<0.0001** |
| Residual | 0.1293 | 228 | 0.0005672 |  |  |

| **Test Details** | **Mean 1** | **Mean 2** | **Mean Diff.** | **SE of Diff** | **N1** | **N2** | **t** | **DF** | **95% Cl of diff** | **Adjusted p-value** |
| --- | --- | --- | --- | --- | --- | --- | --- | --- | --- | --- |
| CD – WD |  |  |  |  |  |  |  |  |  |  |
| Week 0-1 | 0.6629 | 0.7042 | -0.04128 | 0.01196 | 38 | 40 | 3.452 | 75.06 | -0.07179 to -0.01076 | 0.0037 |
| Week 1-2 | 0.682 | 0.6551 | 0.02698 | 0.00921 | 38 | 40 | 2.929 | 71.64 | 0.003442 to 0.05051 | 0.0181 |
| Week 2-3 | 0.6414 | 0.6039 | 0.03747 | 0.00934 | 38 | 40 | 4.008 | 75.77 | 0.01362 to 0.06132 | 0.0006 |
| **Week 3-4** | 0.5689 | 0.5272 | 0.04175 | 0.00724 | 38 | 40 | 5.76 | 73.89 | **0.02324 to 0.06025** | **<0.0001** |

| **ANOVA Table** | **SS** | **DF** | **MS** | **F (DFn, DFd)** | **Adjusted p-value** |
| --- | --- | --- | --- | --- | --- |
| Time x Group | 0.04998 | 15 | 0.003332 | F (15, 300) = 13.04 | P<0.0001 |
| Time | 0.8928 | 5 | 0.1786 | F (2.802, 168.1) = 699.1 | P<0.0001 |
| Group | 0.01572 | 3 | 0.00524 | F (3, 60) = 1.877 | P=0.1431 |
| Subject | 0.1675 | 60 | 0.002791 | F (60, 300) = 10.93 | P<0.0001 |
| Residual | 0.07662 | 300 | 0.0002554 |  |  |

| **Test Details** | **Mean 1** | **Mean 2** | **Mean Diff.** | **SE of Diff** | **N1** | **N2** | **q** | **DF** | **95% Cl of diff** | **Adjusted p-value** |
| --- | --- | --- | --- | --- | --- | --- | --- | --- | --- | --- |
| Week 4-5 |  |  |  |  |  |  |  |  |  |  |
| CDU vs. CDE | 0.4884 | 0.4871 | 0.001312 | 0.01034 | 16 | 16 | 0.1796 | 24.81 | -0.02713 to 0.02976 | 0.9992 |
| CDU vs. WDU | 0.4884 | 0.4721 | 0.016310 | 0.01127 | 16 | 16 | 2.048 | 23.16 | -0.01485 to 0.04748 | 0.4837 |
| CDU vs. WDE | 0.4884 | 0.4669 | 0.02144 | 0.01379 | 16 | 16 | 2.198 | 20.22 | -0.01713 to 0.06001 | 0.4256 |
| CDE vs. WDU | 0.4871 | 0.4721 | 0.015 | 0.01326 | 16 | 16 | 1.6 | 29.61 | -0.02108 to 0.05108 | 0.6735 |
| CDE vs. WDE | 0.4871 | 0.4669 | 0.02013 | 0.01546 | 16 | 16 | 1.840 | 26.74 | -0.02222 to 0.06247 | 0.5701 |
| WDU vs. WDE | 0.4721 | 0.4669 | 0.0051 | 0.0161 | 16 | 16 | 0.450 | 28.31 | -0.03881 to 0.04906 | 0.9886 |
| Week 5-6 |  |  |  |  |  |  |  |  |  |  |
| CDU vs. CDE | 0.4459 | 0.4528 | -0.006937 | 0.00767 | 16 | 16 | 1.279 | 24.78 | -0.02805 to 0.01417 | 0.8026 |
| **CDU vs. WDU** | 0.4459 | 0.4198 | 0.02613 | 0.007213 | 16 | 16 | 5.122 | 26.07 | **0.006341 to 0.04591** | **0.0064** |
| CDU vs. WDE | 0.4459 | 0.4258 | 0.02013 | 0.009069 | 16 | 16 | 3.138 | 21.81 | -0.005077 to 0.04533 | 0.1495 |
| **CDE vs. WDU** | 0.4528 | 0.4198 | 0.03306 | 0.008889 | 16 | 16 | 5.26 | 29.78 | **0.008881 to 0.05724** | **0.0044** |
| CDE vs. WDE | 0.4528 | 0.4258 | 0.02706 | 0.01045 | 16 | 16 | 3.662 | 28.68 | -0.001433 to 0.05556 | 0.0673 |
| WDU vs. WDE | 0.4198 | 0.4258 | -0.006000 | 0.01012 | 16 | 16 | 0.8383 | 27.6 | -0.03366 to 0.02166 | 0.9334 |
| Week 6-7 |  |  |  |  |  |  |  |  |  |  |
| CDU vs. CDE | 0.4053 | 0.4265 | -0.02119 | 0.008423 | 16 | 16 | 3.558 | 29.05 | -0.04413 to 0.001757 | 0.0784 |
| **CDU vs. WDU** | 0.4053 | 0.3649 | 0.04044 | 0.009855 | 16 | 16 | 5.803 | 25.84 | **0.01339 to 0.06748** | **0.0019** |
| CDU vs. WDE | 0.4053 | 0.3863 | 0.019 | 0.009295 | 16 | 16 | 2.891 | 27.1 | -0.006430 to 0.04443 | 0.1972 |
| **CDE vs. WDU** | 0.4265 | 0.3649 | 0.06163 | 0.01048 | 16 | 16 | 8.312 | 28.39 | **0.03302 to 0.09023** | **<0.0001** |
| **CDE vs. WDE** | 0.4265 | 0.3863 | 0.04019 | 0.00996 | 16 | 16 | 5.706 | 29.29 | **0.01307 to 0.06731** | **0.0019** |
| WDU vs. WDE | 0.3649 | 0.3863 | -0.02144 | 0.0112 | 16 | 16 | 2.708 | 29.78 | -0.05190 to 0.009022 | 0.2437 |
| Week 7-8 |  |  |  |  |  |  |  |  |  |  |
| CDU vs. CDE | 0.3640 | 0.3759 | -0.01194 | 0.005954 | 16 | 16 | 2.836 | 29.56 | -0.02814 to 0.004265 | 0.2090 |
| CDU vs. WDU | 0.3640 | 0.3761 | -0.01206 | 0.008078 | 16 | 16 | 2.112 | 23.56 | -0.03438 to 0.01025 | 0.4572 |
| **CDU vs. WDE** | 0.3640 | 0.3919 | -0.02788 | 0.006923 | 16 | 16 | 5.694 | 26.72 | **-0.04683 to -0.008916** | **0.0022** |
| CDE vs. WDU | 0.3759 | 0.3761 | -0.000125 | 0.008341 | 16 | 16 | 0.0212 | 25.35 | -0.02305 to 0.02280 | >0.9999 |
| CDE vs. WDE | 0.3759 | 0.3919 | -0.01594 | 0.007228 | 16 | 16 | 3.118 | 28.38 | -0.03566 to 0.003781 | 0.1463 |
| WDU vs. WDE | 0.3761 | 0.3919 | -0.01581 | 0.009058 | 16 | 16 | 2.469 | 28.72 | -0.04051 to 0.008881 | 0.3196 |
| Week 8-9 |  |  |  |  |  |  |  |  |  |  |
| CDU vs. CDE | 0.3416 | 0.3558 | -0.01419 | 0.005475 | 16 | 16 | 3.665 | 29.99 | -0.02907 to 0.0006987 | 0.0661 |
| CDU vs. WDU | 0.3416 | 0.3576 | -0.01606 | 0.007819 | 16 | 16 | 2.905 | 24 | -0.03763 to 0.005507 | 0.1968 |
| **CDU vs. WDE** | 0.3416 | 0.3674 | -0.02588 | 0.006742 | 16 | 16 | 5.427 | 27.1 | **-0.04432 to -0.007429** | **0.0036** |
| CDE vs. WDU | 0.3558 | 0.3576 | -0.001875 | 0.00778 | 16 | 16 | 0.3408 | 23.71 | -0.02336 to 0.01961 | 0.9949 |
| CDE vs. WDE | 0.3558 | 0.3674 | -0.01169 | 0.006697 | 16 | 16 | 2.468 | 26.80 | -0.03002 to 0.006647 | 0.3211 |
| WDU vs. WDE | 0.3576 | 0.3674 | -0.0098 | 0.008719 | 16 | 16 | 1.592 | 28.78 | -0.03358 to 0.01395 | 0.6771 |
| Week 9-10 |  |  |  |  |  |  |  |  |  |  |
| CDU vs. CDE | 0.3258 | 0.3376 | -0.01181 | 0.005209 | 16 | 16 | 3.207 | 29.97 | -0.02598 to 0.002351 | 0.1285 |
| CDU vs. WDU | 0.3258 | 0.3376 | -0.01188 | 0.006188 | 16 | 16 | 2.714 | 27.99 | -0.02877 to 0.005021 | 0.2433 |
| **CDU vs. WDE** | 0.3258 | 0.3454 | -0.01969 | 0.005698 | 16 | 16 | 4.886 | 29.45 | **-0.03520 to -0.004177** | **0.0087** |
| CDE vs. WDU | 0.3376 | 0.3376 | -0.000063 | 0.006114 | 16 | 16 | 0.0145 | 27.54 | -0.01677 to 0.01665 | >0.9999 |
| CDE vs. WDE | 0.3376 | 0.3454 | -0.007875 | 0.00562 | 16 | 16 | 1.982 | 29.16 | -0.02318 to 0.007425 | 0.5083 |
| WDU vs. WDE | 0.3376 | 0.3454 | -0.00781 | 0.006536 | 16 | 16 | 1.69 | 29.45 | -0.02560 to 0.009980 | 0.6346 |

**Supplemental Table 4.** Weekly weight-corrected calorie intake. Caloric intake values were calculated by multiplying the total food consumption per animal by the total kcal (S Table 1) for each respective diet. Two-way repeated-measures ANOVA was performed, with diet and time as factors, during the first four weeks. A TW-RM ANOVA was used to evaluate the data from week 5 after groups underwent exposure to psychosocial stress to the end of the study, with time and treatment (diet + stress) as factors.

| **ANOVA table** | **SS** | **DF** | **MS** | **F (DFn, DFd)** | **Adjusted p-value** |
| --- | --- | --- | --- | --- | --- |
| Time x Diet | 2.227 | 3 | 0.7425 | F (3, 228) = 75.16 | P<0.0001 |
| Time | 15.69 | 3 | 5.229 | F(2.052, 156.0) = 529.3 | P<0.0002 |
| Diet | 14.55 | 1 | 14.55 | F (1, 76) = 148.1 | P<0.0003 |
| Subject | 7.469 | 76 | 0.09828 | F (76, 228) 9.949 | P<0.0004 |
| Residual | 2.252 | 228 | 0.009879 |  |  |

| **Test Details** | **Mean 1** | **Mean 2** | **Mean Diff** | **SE of Diff.** | **N1** | **N2** | **t** | **DF** | **95% CI of Diff.** | **Adjusted p-value** |
| --- | --- | --- | --- | --- | --- | --- | --- | --- | --- | --- |
| Week 4-5 |  |  |  |  |  |  |  |  |  |  |
| CDU vs. CDE | 1.856 | 1.851 | 0.01 | 0.0395 | 16 | 16 | 0.127 | 24.91 | -0.1367 to 0.1467 | >0.9999 |
| **CDU vs. WDU** | 1.856 | 2.171 | -0.32 | 0.0499 | 16 | 16 | 6.317 | 20.97 | **-0.4981 to -0.1319** | **0.0001***** |
| **CDU vs. WDE** | 1.856 | 2.148 | -0.29 | 0.0622 | 16 | 16 | 4.682 | 18.67 | **-0.5239 to -0.05863** | **0.0061**** |
| **CDE vs. WDU** | 1.851 | 2.171 | -0.32 | 0.0565 | 16 | 16 | 5.664 | 27.67 | **-0.5202 to -0.1198** | **0.0002***** |
| **CDE vs. WDE** | 1.851 | 2.148 | -0.27 | 0.0676 | 16 | 16 | 4.3800 | 23.91 | **-0.5401 to -0.05238** | **0.0073**** |
| WDU vs. WDE | 2.171 | 2.148 | 0.02 | 0.0742 | 16 | 16 | 0.320 | 28.22 | -0.2385 to 0.2860 | >0.9999 |
| Week 5-6 |  |  |  |  |  |  |  |  |  |  |
| CDU vs. CDE | 1.693 | 1.721 | -0.03 | 0.0289 | 16 | 16 | 0.974 | 24.62 | -0.1319 to 0.07566 | >0.9999 |
| **CDU vs. WDU** | 1.693 | 1.931 | -0.24 | 0.0312 | 16 | 16 | 7.641 | 23.18 | **-0.3510 to -0.1253** | **<0.0001****** |
| **CDU vs. WDE** | 1.693 | 1.959 | -0.27 | 0.0403 | 16 | 16 | 6.597 | 19.65 | **-0.4149 to -0.1163** | **<0.0001****** |
| **CDE vs. WDU** | 1.721 | 1.931 | -0.21 | 0.0369 | 16 | 16 | 5.691 | 29.7 | **0.3398 to -0.08022** | **0.0001***** |
| **CDE vs. WDE** | 1.721 | 1.959 | -0.24 | 0.0449 | 16 | 16 | 5.2960 | 26.02 | **-0.3975 to -0.07746** | **0.0006***** |
| WDU vs. WDE | 1.931 | 1.959 | -0.03 | 0.0464 | 16 | 16 | 0.593 | 27.48 | -0.1919 to 0.1369 | >0.9999 |
| Week 6-7 |  |  |  |  |  |  |  |  |  |  |
| CDU vs. CDE | 1.656 | 1.741 | -0.09 | 0.0373 | 16 | 16 | 2.297 | 29.26 | -0.2170 to 0.04571 | 0.6531 |
| CDU vs. WDU | 1.656 | 1.491 | 0.17 | 0.0429 | 16 | 16 | 3.850 | 26.49 | **0.01237 to 0.3176** | **0.0240*** |
| CDU vs. WDE | 1.656 | 1.578 | 0.08 | 0.0405 | 16 | 16 | 1.912 | 27.69 | -0.06611 to 0.2211 | 0.9152 |
| **CDE vs. WDU** | 1.741 | 1.491 | 0.25 | 0.0454 | 16 | 16 | 5.524 | 28.65 | **0.09044 to 0.4108** | **0.0002***** |
| **CDE vs. WDE** | 1.741 | 1.578 | 0.16 | 0.0432 | 16 | 16 | 3.7780 | 29.46 | **0.01113 to 0.3151** | **0.0254*** |
| WDU vs. WDE | 1.491 | 1.578 | -0.09 | 0.0481 | 16 | 16 | 1.820 | 29.79 | -0.2566 to 0.08155 | 0.9480 |
| Week 7-8 |  |  |  |  |  |  |  |  |  |  |
| CDU vs. CDE | 1.436 | 1.486 | -0.05 | 0.0238 | 16 | 16 | 2.099 | 29.47 | -0.1339 to 0.03385 | 0.8057 |
| CDU vs. WDU | 1.436 | 1.481 | -0.05 | 0.0320 | 16 | 16 | 1.428 | 23.65 | -0.1610 to 0.06973 | 0.9986 |
| **CDU vs. WDE** | 1.436 | 1.543 | -0.11 | 0.0275 | 16 | 16 | 3.893 | 26.76 | **-0.2045 to -0.009225** | **0.0212*** |
| CDE vs. WDU | 1.486 | 1.481 | 0.00 | 0.0331 | 16 | 16 | 0.132 | 25.62 | -0.1140 to 0.1228 | >0.9999 |
| CDE vs. WDE | 1.486 | 1.543 | -0.06 | 0.0288 | 16 | 16 | 1.9750 | 28.56 | -0.1585 to 0.04484 | 0.8838 |
| WDU vs. WDE | 1.481 | 1.543 | -0.06 | 0.0358 | 16 | 16 | 1.710 | 28.75 | -0.1876 to 0.06515 | 0.9756 |
| Week 8-9 |  |  |  |  |  |  |  |  |  |  |
| CDU vs. CDE | 1.349 | 1.401 | -0.05 | 0.0216 | 16 | 16 | 2.407 | 30 | -0.1276 to 0.02385 | 0.5583 |
| CDU vs. WDU | 1.349 | 1.411 | -0.06 | 0.0305 | 16 | 16 | 2.009 | 23.98 | -0.1712 to 0.04866 | 0.8743 |
| **CDU vs. WDE** | 1.349 | 1.448 | -0.10 | 0.0271 | 16 | 16 | 3.643 | 26.42 | **-0.1953 to -0.002193** | **0.0408*** |
| CDE vs. WDU | 1.401 | 1.411 | -0.01 | 0.0305 | 16 | 16 | 0.307 | 23.99 | -0.1193 to 0.1005 | >0.9999 |
| CDE vs. WDE | 1.401 | 1.448 | -0.05 | 0.0271 | 16 | 16 | 1.7290 | 26.43 | -0.1434 to 0.04970 | 0.9730 |
| WDU vs. WDE | 1.411 | 1.448 | -0.04 | 0.0347 | 16 | 16 | 1.082 | 29.23 | -0.1596 to 0.08457 | >0.9999 |
| Week 9-10 |  |  |  |  |  |  |  |  |  |  |
| CDU vs. CDE | 1.286 | 1.33 | -0.04 | 0.0212 | 16 | 16 | 2.092 | 30 | -0.1189 to 0.03016 | 0.8095 |
| CDU vs. WDU | 1.286 | 1.333 | -0.05 | 0.0246 | 16 | 16 | 1.930 | 28.22 | -0.1345 to 0.03952 | 0.9064 |
| CDU vs. WDE | 1.286 | 1.363 | -0.08 | 0.0227 | 16 | 16 | 3.389 | 29.58 | -0.1567 to 0.002944 | 0.0697 |
| CDE vs. WDU | 1.330 | 1.333 | 0.00 | 0.0245 | 16 | 16 | 0.127 | 28.12 | -0.08994 to 0.08369 | >0.9999 |
| CDE vs. WDE | 1.330 | 1.363 | -0.03 | 0.0226 | 16 | 16 | 1.4370 | 29.53 | -0.1121 to 0.04708 | 0.9982 |
| WDU vs. WDE | 1.333 | 1.363 | -0.03 | 0.0258 | 16 | 16 | 1.138 | 29.45 | -0.1203 to 0.06153 | >0.9999 |
